# Decomposing parasite fitness in a two-host, two-parasite system reveals the underpinnings of parasite specialization

**DOI:** 10.1101/256974

**Authors:** Eva J. P. Lievens, Julie Perreau, Philip Agnew, Yannis Michalakis, Thomas Lenormand

## Abstract

The ecological specialization of parasites – whether they can obtain high fitness on very few or very many different host species – is a determining feature of their ecology. In order to properly assess specialization, it is imperative to measure parasite fitness across host species; to understand its origins, fitness must be decomposed into the underlying traits. Despite the omnipresence of parasites with multiple hosts, very few studies assess and decompose their specialization in this way. To bridge this gap, we quantified the infectivity, virulence, and transmission rate of two parasites, the horizontally transmitted microsporidians *Anostracospora rigaudi* and *Enterocytospora artemiae*, in their natural hosts, the brine shrimp *Artemia parthenogenetica* and *Artemia franciscana*. Our results demonstrate that each parasite performs well on one of the two host species (*A. rigaudi* on *A. parthenogenetica*, and *E. artemiae* on *A. franciscana*), and poorly on the other. This partial specialization is driven by high infectivity and transmission rates in the preferred host, and is associated with maladaptive virulence and large costs of resistance in the other. Our study represents a rare empirical contribution to the study of parasite evolution in multi-host systems, highlighting the negative effects of under- and over-exploitation when adapting to multiple hosts.

## Introduction

Life in a variable environment imposes an evolutionary choice between specializing to certain habitats and remaining a generalist. This dilemma is particularly pressing for parasitic species, which often come into contact with a wide range of potential habitats (i.e. hosts). Evolving an optimal level of specialization is not trivial, as adaptation to one host may come at the expense of adaptation to another (Levins 1968, Kawecki 1994, Kassen 2002). Furthermore, the degree of specialization affects the ecology and future evolution of the parasite: generalist parasites are more likely to survive perturbations in the host community and to colonize new hosts (Cleaveland et al. 2001, Agosta et al. 2010), while specialist parasites are more likely to interact tightly with their hosts (Kawecki 1998). The degree of specialization, therefore, is a key trait of parasite species. It varies widely among species – even within clades, parasites can range from extremely specific (infecting only one host species) to widely generalist (infecting tens of host species) (Poulin and Keeney 2008) – and through time – many parasites can evolve from generalism to specialism or vice-versa when conditions change (e.g. Desdevises et al. 2002, Tanaka et al. 2007, Johnson et al. 2009, Cenzer 2016).

Thus, assessing how specialized multi-host parasites are, and to which hosts, is an essential step to understanding and controlling the epidemiology and evolution of multi-host parasites. To this end, the ‘ecological specialization’ of parasites should be distinguished from the standard concepts ‘host range’ and ‘host specificity’ (sensu Lymbery 1989). Neither host range – the number of host species in which a parasite occurs – nor host specificity – host range weighted by infection intensity or host phylogeny – account for the existence of host species that barely contribute to the parasite’s transmission. Such ‘spillover’ hosts (sensu Fenton et al. 2015) can readily become infected, but do not transmit the parasite enough to keep its population growth rate above one. As a consequence, infection in the spillover hosts quickly dies out if there is no replenishing transmission from suitable hosts (“dead-end” and “stuttering chain” dynamics, Viana et al. 2014). In essence, these are ecological source-sink dynamics. Ecological specialization can take these dynamics into account: it is based on niche breadth (Futuyma and Moreno 1988), and sink habitats fall outside the fundamental niche (Pulliam 1988). Classifying organisms as ecological generalists or specialists means studying the variation in their fitness across a range of environments (Kassen 2002). Applied to parasites, this means their fitness must be assessed in all the affected host species. Such assessments typically require detailed epidemiological models (e.g. Rhodes et al. 1998, Fenton et al. 2015) or sizeable experiments (Jaenike and Dombeck 1998, Ahonen et al. 2006, Auld et al. 2017).

A second step is to understand why parasite fitness varies across hosts. The fitness of infections emerges from a suite of parasite- and host-determined traits, including infectivity, exploitation of host resources, virulence, immune evasion, and transmission success. The nature of these traits has important consequences for a parasite: evolutionary constraints can emerge from functional correlations between traits within a host species (Walther and Ewald 2004, Alizon et al. 2009, Alizon and Michalakis 2015, Hall et al. 2017), or from correlations between the same trait in different host species (Futuyma and Moreno 1988, Via and Hawthorne 2002). They also determine the source of the parasite’s maladaptation to spillover hosts (Woolhouse et al. 2001). This has been best studied with regards to virulence and transmission, mostly in single-host systems (e.g. Dwyer et al. 1990, Fraser et al. 2007, de Roode et al. 2008, Doumayrou et al. 2012). Studies that decompose the fitness of multi-host parasites into component traits are very rare (reviewed in Rigaud et al. 2010)(Agudelo-Romero et al. 2008, Auld et al. 2017).

Here, we examine specialization and its component traits in a natural multi-host, multi-parasite system. In the saltern of Aigues-Mortes, France, two species of brine shrimp occur in sympatry: a native parthenogenetic clade, *Artemia parthenogenetica*, and an introduced sexual species, *Artemia franciscana* (Amat et al. 2005). Both *Artemia* species are parasitized by the microsporidians *Anostracospora rigaudi* and *Enterocytospora artemiae*. These parasites infect the gut epithelium, transmitting infection horizontally through spores released with the faeces (Rode et al. 2013b, 2013a). Since the saltern lacks spatial structure (Nougué et al. 2015), the pool of microsporidian spores is shared between *A. franciscana* and *A. parthenogenetica* (cf. Fels 2006). Although the rates of inter-specific transmission should therefore be high, and both *A. rigaudi* and *E. artemiae* commonly infect either host species, the two microsporidians appear to be somewhat specialized: *A. rigaudi* is always more prevalent in *A. parthenogenetica*, and is also dependent on this species to maintain itself in the host community; in contrast, *E. artemiae* is consistently more infectious to and more prevalent in *A. franciscana* (Lievens et al. subm.). Historically, the association of *A. parthenogenetica* and *A. rigaudi* predates the introduction of *A. franciscana* (in 1970, Rode et al. 2013c), while *A. franciscana* is also infected by *E. artemiae* in its native range (Rode et al. 2013c). It is not known whether *E. artemiae* was also present in France before the introduction of *A. franciscana*, whether it was co-introduced, or whether it arrived independently afterwards.

We evaluated parasite specialization in this system by studying the infectivity, virulence, and transmission of *A. rigaudi* and *E. artemiae* in each of their hosts. We confirm experimentally that while both microsporidians can complete their life cycle in the two host species, neither is a complete generalist. Rather, *A. rigaudi* is largely specialized on *A. parthenogenetica*, while *E. artemiae* is largely specialized on *A. franciscana*. Further, we show that the lower fitness of the two parasites in their non-specialized hosts was caused by a reduction in infectivity and transmission rate (in both cases), combined with a suboptimal degree of virulence (too low for *E. artemiae*; too high for *A. rigaudi*). This demonstrates that a successful calibration of host exploitation and parasite virulence is central to the specialization of multi-host parasites.

## Methods

We performed two experiments to investigate the life history and virulence of the microsporidians *A. rigaudi* and *E. artemiae* in their *Artemia* hosts. First, we used dose-response tests to quantify infectivity in each host-parasite combination. Second, we did a large-scale experimental infection experiment, tracking host growth, mortality, and reproduction, as well as parasite transmission, over a period of two months. These results allowed us to estimate the virulence and fitness of each parasite on each host.

### Experimental conditions

The *Artemia* used in both experiments were raised in the lab in parasite-free conditions. *A. franciscana* were hatched from dormant cysts sampled from the saltern of Aigues-Mortes, France, and stored in dry conditions at 4 °C. We used three batches of cysts, sampled at the sites Caitive Nord or Caitive Sud in October 2013 or 2014. Cysts were hatched following the protocol described by Lievens et al. (2016). *A. parthenogenetica* were collected as live larvae from a mix of clones. The *A. parthenogenetica* clones were started by females collected in Aigues-Mortes, who were allowed to multiply and produce cysts in the lab; those cysts were then hatched to produce parasite-free stock lines. All *Artemia* were maintained at 23 ± 1 °C, in a parasite-free 90 ppt saline medium produced by diluting concentrated, autoclaved brine (Camargue Pêche, France) with deionized water. *Artemia* were fed *ad libitum* with freeze-dried microalgae (*Tetraselmis chuii*, Fitoplankton marino, Spain) dissolved in deionized water. Experimental conditions matched the cultivation conditions, except that feeding was regulated (see below).

We created stocks of *A. rigaudi* and *E. artemiae* for use in the experiment by combining infected *Artemia* from various sites in Aigues-Mortes between October 2014 and March 2015. We added new infected hosts to the stocks whenever we found field populations that were heavily infected with either *A. rigaudi* or *E. artemiae*. We also regularly added uninfected, lab-bred *Artemia* to help maintain the infection. We selected both infected *A. franciscana* and infected *A. parthenogenetica* from the field, and maintained each stock population on a mix of *A. franciscana* and *A. parthenogenetica* hosts (n_hosts_ at any given time=∼20-∼50 per microsporidian species). Thus, our stocks contained a mix of spores from different field sites and times, collected from and propagated on both host species.

### Spore collection and quantification

To produce the inocula for Experiments 1 and 2, we collected spores from the lab stocks of *A. rigaudi* and *E. artemiae* described above. The stock hosts were kept in large separating funnels, so that their feces (containing spores) settled down into the funnel’s tube and could be collected easily. For our experiments, we collected feces produced over 20-hour periods (feces suspended in ∼15 mL). Because fecal aggregates can trap spores and skew concentration estimates, we homogenized the fecal solutions by dividing them into 1.2 mL Qiagen Collection Microtubes, adding two 4 mm stainless steel beads to each tube, and shaking them at 30 Hz for 30 s. Once homogenized, the fecal solutions were recombined to their original volume. To quantify the spore concentration in the fecal solutions, we took 1 mL subsamples and added 10 µL 1X Calcofluor White Stain (18909 Sigma-Aldrich, USA) to each. After staining for 10 min, we rinsed the subsamples by centrifuging them for 8 min at 10000 g, replacing 910 µL of the supernate with 900 µL deionized water and vortexing well.

We then concentrated the subsamples to 20X by repeating the centrifugation step and removing 950 µL of the supernate. Finally, we estimated the concentration by counting the number of spores in 0.1 µL on a Quick Read counting slide (Dominique Dutscher) under a Zeiss AX10 fluorescence microscope (10x 40x magnification; excitation at 365 nm; Zeiss filter set: 62 HE BFP + GFP + HcRed shift free (E)). We repeated the counts twice (for Experiment 2) or thrice (for Experiment 1); the spore concentration per µL in the unconcentrated fecal solutions was then equal to (mean of the spore counts*10)/20. Finally, we added 90 ppt clean saline medium to the homogenized fecal solutions until the correct concentration for inoculation was reached.

### Experiment 1: Infectivity

#### Experimental design and execution

Previous papers studied the infectivity of *A. rigaudi* and *E. artemiae* using single, uncontrolled spore doses (Lievens et al. subm., Rode et al. 2013a). Here, we quantified infectivity more precisely by exposing individual *A. parthenogenetica* and *A. franciscana* to a range of controlled spore doses and measuring the proportion of infected individuals.

We exposed experimental hosts to doses of 0, 400, 800, 1600, 3200 and 6400 spores per individual. To ensure hosts ingested all spores, each host was first exposed in a highly-concentrated medium: individuals were placed in 2 mL Eppendorf tubes with 0.45 mL spore solution, 1 mL extra brine and 0.25 mL algal solution (3.4*10^9^ *T. chuii* cells/L deionized water). After two days, hosts were transferred to 40 mL glasses containing 20 mL brine and the infection was allowed to incubate for three more days; hosts were fed a total of 1 mL algal solution over the three days. Surviving hosts were then sacrificed and tested for the presence of *A. rigaudi* or *E. artemiae* by PCR (following Rode et al. 2013a). Treatments were replicated 20 times, except when spore availability was limiting (*E. rigaudi* on *A. parthenogenetica*: 16, 8 and 4 replicates for the doses 400, 3200 and 6400 spores per individual, respectively). All hosts were ∼4 weeks old and measured between 5 and 8 mm; *A. franciscana* hosts were mixed males and females.

#### Statistical analyses

To analyze the dose-response curves, we used four-parameter log-logistic modeling in R (package drc, Ritz and Strebig 2005, R Core Team 2014). In these models, the four parameters determining the shape of the sigmoidal curve are: the lower limit (set to 0 in our case), the upper limit, the slope around the point of inflection, and the point of inflection (which here is the same as the ED_50_). The (binomial) response variable was the number of individuals that were infected vs. uninfected. Because we did not perform the *A. parthenogenetica* and *A. franciscana* experiments at the same time, we could not control for environmental effects. Thus, we simply tested if the dose-response curves for *A. rigaudi* and *E. artemiae* were different within each host species. To do this, we fit models that did or did not include a ‘microsporidian species’ effect and compared the two using a likelihood ratio test. If the effect was significant, we went on to compare the parameters of the two resulting curves (‘compParm’ function in the drc package).

### Experiment 2: Virulence and transmission

#### Experimental design and execution

To quantify the virulence and transmission rates of *A. rigaudi* and *E. artemiae*, we experimentally infected individual *Artemia* with controlled spore doses. We then tracked their survival, growth, reproductive output, and spore production over a two-month period. We also quantified host-to-host transmission at two time points.

*A. franciscana* males, *A. franciscana* females, and *A. parthenogenetica* females were divided into three treatments: ‘Controls’, ‘Exposure to *A. rigaudi*, and ‘Exposure to *E. artemiae*’, which were replicated as permitted by spore and host availability (Table 1). *A. franciscana* hosts were subdivided into three blocks, determined by their origin: Caitive Nord 2013, Caitive Nord 2014, or Caitive Sud 2014. *A. parthenogenetica* hosts were subdivided into two blocks, determined by the age of their batch: 34 ± 2 or 26 ± 2 days (because the relative contribution of the different clones to the batches was not controlled, the genotype frequencies of these groups could differ). All hosts were subadults (adult body plan but sexually immature). All *A. franciscana* were aged 38 ± 1 days and measured 4.5 or 5.0 mm; *A. parthenogenetica* measured 6.5, 7.0 or 7.5 mm. Size classes were evenly distributed across blocks and treatments.

We exposed experimental hosts to spore doses designed to be comparable while maximizing infection rate (see results of Experiment 1): 3000 spores/individual for *A. rigaudi* and 2500 spores/individual for *E. artemiae*. Because *A. parthenogenetica* had low infection rates with *E. artemiae*, a separate set of *A. parthenogenetica* was infected with 10000 *E. artemiae* spores per individual (Table 1). To ensure hosts ingested all spores, each host was exposed in a highly-concentrated medium over a two-day period: individuals were placed in 2 mL Eppendorf tubes with 0.37 mL spore solution and 1.25 mL brine containing 2.6*10^6^ *T. chuii* cells.

**Table 1.** Number of replicates for the different treatments and blocks. See ‘Methods – Experiment 2’ for more information.

| Treatment:<br>[spore dose] | Exposure to *A. rigaudi*<br>[3 000 sp/i] |  | Exposure to *E. artemiae*<br>[2 500 sp/i] |  | Exposure to *E. artemiae*<br>[10 000 sp/i] |  | Controls |  |
| --- | --- | --- | --- | --- | --- | --- | --- | --- |
| ***A. franciscana*** | **86 ♂** | **86 ♀** | **132 ♂** | **132 ♀** |  |  | **120 ♂** | **120 ♀** |
| Origin: Caitive Nord 2013 | 26 ♂ | 26 ♀ | 72 ♂ | 72 ♀ |  |  | 60 ♂ | 60 ♀ |
| Origin: Caitive Nord 2014 | 30 ♂ | 30 ♀ | 30 ♂ | 30 ♀ |  |  | 30 ♂ | 30 ♀ |
| Origin: Caitive Sud 2014 | 30 ♂ | 30 ♀ | 30 ♂ | 30 ♀ |  |  | 30 ♂ | 30 ♀ |
| ***A. parthenogenetica*** |  | **96 ♀** |  | **96 ♀** |  | **33 ♀** |  | **96 ♀** |
| Batch: 34 ± 2 days old |  | 48 ♀ |  | 48 ♀ |  | 18 ♀ |  | 48 ♀ |
| Batch: 26 ± 2 days old |  | 48 ♀ |  | 48 ♀ |  | 15 ♀ |  | 48 ♀ |

After exposure (= on day 1 of the experiment), individuals were transferred to open tubes, which rested upright in 40 mL plastic cups containing 20 mL of brine. The lower end of the tube was fitted with a 1×1 mm net. The netting prevented experimental (adult) individuals from swimming to the bottom of the cup, while allowing spores, feces, and offspring to pass through; this limited secondary infections from a host’s own feces. Cups were randomly placed in trays, which were routinely rotated to standardize effects of room placement. Water was changed every five days. Hosts were fed 0.5 mL algal solution daily (2.6*10^9^ *T. chuii* cells/L deionized water); this feeding regime corresponds to half of the maximum ingestion rate of an adult *Artemia* (Reeve 1963) and has been shown to reveal energetic trade-offs (Rode et al. 2011). We ended our experiment after 60 days, at which point surviving individuals were sacrificed and tested for infection by PCR (following Rode et al. 2013a).

To quantify the effects of infection on the hosts, we tracked the growth, survival, and reproduction of the experimental individuals. Body length was recorded on days 30 and 60. Survival was recorded daily; dead individuals were tested for infection by PCR (following Rode et al. 2013a). We did not track reproduction for males, because male reproductive success is heavily influenced by the female partner (e.g. female clutch size). For females, measures of reproductive success were recorded daily, including date of sexual maturity (first detection of a fully-formed ovisac or of yolk accumulation in oocytes, Metalli and Ballardin 1970), clutch date, clutch type, and clutch size. *Artemia* females are iteroparous, producing on average one clutch per five days (Bowen 1962, Metalli and Ballardin 1970). Clutches may be of two types: live larvae (‘nauplii’), or dormant encysted embryos (‘cysts’). Neonatal nauplii are barely visible to the eye and have high death rates. For ease of measurement, therefore, clutches of nauplii were counted five days after sighting. During these five days nauplii were in competition for resources with their mother (plus an additional male if *A. franciscana*, see below). However, mothers were removed at each water change, which could happen before the clutch had reached the five-day mark. In these cases, we placed a new tube containing one (or two, if *A. franciscana*) adult male *Artemia* above the nauplii to ensure the same level of food competition.

While *A. parthenogenetica* females reproduce in isolation, *A. franciscana* females need to be fertilized before each clutch (Bowen 1962). We therefore added mature *A. franciscana* males from parasite-free lab stocks to each tube containing an *A. franciscana* female. To prevent cross-contamination between the male and the female, exposed males were removed and new uninfected males added every five days (five-day estimate based on infection detection time as found by Rode et al. 2013a). Male *Artemia* mate-guard by clasping females around the abdomen (Bowen 1962), and forcible removal may be harmful to both partners. To avoid this, males found mate-guarding on the fifth day were given up to two extra days with the female, after which they were forcibly removed. Couples were fed twice the individual food allocation.

To estimate parasite fitness, we estimated spore production at regular points throughout the experiment. To do this, we collected 1 mL of feces (containing parasite spores) from every experimentally infected host at every water change. Samples were stored in 1.2 mL Qiagen Collection Microtubes and refrigerated until the spore concentration could be quantified. To measure the spore concentration, we homogenized and stained each sample as described above, with minor differences in the centrifugation steps (16 min at 5000 g) and the final concentration (concentrated to 14.3X by removing 930 µL of the supernate). Spores were counted once per sample, as described above. Because counting spores is labor-intensive, we restricted our efforts to the feces samples collected on days 15, 30, 45 and 60.

We also investigated the host-to-host transmission success of the parasites and its relation to spore production. On days 30 and 60, we allowed a subset of experimental hosts (hereafter the ‘donors’) to infect groups of uninfected ‘recipient’ hosts for 24 hours. Donors were first placed with either eight *A. franciscana* or eight *A. parthenogenetica* recipients; after 24 hours, the donor was removed and placed with a new group of recipients of the other species. All recipient hosts were taken at random from uninfected lab stocks of varying ages and sizes (min=4 mm, max=10 mm). The donor host was separated from the recipients by a 1×1 mm net; recipients swam underneath them in 40 mL of brine. Infection was allowed to incubate in the recipients for six days after the donor was removed; surviving recipients were then sacrificed and PCR-tested for infection (following Rode et al. 2013a). The prevalence of infection in recipient individuals could then be compared to the number of spores counted in the feces samples on day 30 or 60.

A key aspect of infection follow-up experiments is knowing which individuals were infected after exposure to the parasite, and which were not. In our experiment, testing by PCR was often not sufficient to determine if an individual was infected, because individuals that died before day 60 often had quickly decaying corpses and thus degraded DNA. We therefore considered that an individual was infected if it tested positive by PCR or produced spores or transmitted the infection to a recipient host. If none of these requirements were met, we considered that the individual was not infected. By applying these criteria, we could be sure of the infection status for almost all individuals that died on or after day 15 (the first spore collection date); for any individuals who died before day 15 and who tested negative by PCR, we could not exclude the possibility that they were infected.

#### Statistical analyses: virulence & transmission

We analyzed the results of this experiment in two major parts. First, we examined the virulence of infections (effect of the parasite on host survival, growth, reproduction, and overall fitness). In these analyses, we excluded all individuals that did not become infected after exposure to the parasite. We also excluded all individuals that died before day 15 (we could not be certain of infection status before this day, see above). To make sure that we were not missing important events occurring before this cutoff, we repeated all statistical models for exposed vs. control individuals that died before day 15. Second, we analyzed parasite transmission (spore production rate, infectiousness, and overall fitness). An overview of the analyses is given in Table 2; a detailed description can be found in the Supplementary Material. Below, we describe only those response variables that are not intuitive.

**Table 2.** Overview of statistical analyses. See Supplementary Material for details.

| Tested variable | Statistical models and tests | Fixed-effect terms in the full model | Random/Fraily terms |
| --- | --- | --- | --- |
| **Virulence of infections: *A. franciscana* and *A. parthenogenetica* analyzed separately** |  |  |  |
| **Survival** | Survival models <sup>1</sup> + LRT + Dunnett p.-h. | Treatment <sup>2</sup>, Sex (*A. f.*), Size class, double interactions | Origin (*A. f.*), Batch (*A. p.*) |
| **Growth between days 1 & 30** <sup>3</sup> | LMM + LRT + Dunnett p.-h. | Treatment <sup>2</sup> * Sex (*A. f.*) * Size class | Origin (*A. f.*), Batch (*A. p.*) |
| **Reproduction** |  |  |  |
| Time until sexual maturity | Survival models <sup>1</sup> + LRT + Dunnett p.-h. | Treatment <sup>2</sup> * Size class | Origin (*A. f.*), Batch (*A. p.*) |
| Probability of producing a clutch | Bernouilli GLMM + LRT + Dunnett p.-h. | Treatment <sup>2</sup> * Size class | Origin (*A. f.*), Batch (*A. p.*) |
| Rate of offspring production <sup>4,5,a</sup> | LMM + LRT + Dunnett p.-h. | Treatment <sup>2</sup> * Size class | Origin (*A. f.*), Batch (*A. p.*) |
| Timing of offspring production <sup>4,6,a</sup> | Neg. binomial GLMM + LRT + Dunnett p.-h. | Treatment <sup>2</sup> * (Elapsed % of reproductive period + Elapsed % of reproductive period<sup>2</sup>) | Individual, Origin (*A. f.*), Batch (*A. p.*) |
| Type of offspring produced <sup>a</sup> | Binomial GLMM + LRT + Dunnett p.-h. | Treatment <sup>2</sup> * Size class | Origin (*A. f.*), Batch (*A. p.*) |
| **Fitness (Lifetime reproductive success)** <sup>4,7</sup> | Neg. binomial hurdle models + LRT + Dunnett p.-h. | Treatment <sup>2</sup> * Size class | NA |
| **Parasite transmission and fitness: infections of *A. franciscana* and *A. parthenogenetica* analyzed together** |  |  |  |
| **Infectiousness of one spore, *p*** | LMM + LRT + Tukey p.-h. | Recipient sp. * Parasite sp. | Individual |
| **Spore production rate** <sup>b</sup> |  |  |  |
| Spore count <sup>8,c</sup> | Neg. binomial GLMM + LRT + Tukey p.-h. | Host sp. * Parasite sp. | Individual |
| Spore count ~ dose <sup>8,d</sup> | Neg. binomial GLMM + LRT | Dose | Individual |
| **Fitness** <sup>b</sup> |  |  |  |
| Lifetime transmission success | Kruskal-Wallis tests + Dunn p.-h. | Host-parasite combination <sup>2</sup> | NA |
| Asymptotic growth rate | Kruskal-Wallis tests + Dunn p.-h. | Host-parasite combination <sup>2</sup> | NA |

**Notes:** <sup>1</sup>Survival models were parametric; the best survival distribution was chosen by AICc. <sup>2</sup>*A. parthenogenetica* exposed to low and high doses of *E. artemiae* treated separately. <sup>3</sup>Most host growth occurred between days 1 and 30 (Supp. Table 2), so only this period was analyzed further. <sup>4</sup>Offspring could be nauplii or cysts. These two offspring types were not directly comparable: they probably require different amounts of energy to produce, and we allowed mortality to occur before counting nauplii. To account for this, we repeated the tests with nauplii weighted twice, equally, or half as much as cysts, and based our conclusions on the overall pattern. <sup>5</sup>Rate of offspring production = total number of offspring / length of the reproductive period. The length of the reproductive period was the difference between the date of death (or censoring) and the date of sexual maturity. <sup>5</sup>Modeled as clutch size as a function of the elapsed proportion of the reproductive period. The reproductive period started at sexual maturity and ended at death (or censoring). <sup>7</sup>LRS calculated as the total number of offspring produced over the study period. <sup>8</sup>Spore count = the number of spores counted in the fecal sample; we did not transform the spore count to spores/mL ( ≈ spore count \* 700) to avoid skewing the error distribution.
**Subsets:** <sup>a</sup> Only for females that produced at least 1 clutch. <sup>b</sup> Analyzed for infected individuals only. <sup>c</sup> Excluded *A. p.* exposed to high doses of *E. artemiae*. <sup>d</sup> Only for *A. p.* infected with *E. artemiae*.
**Abbreviations:** GLMM, generalized linear mixed models. LMM, linear mixed models. LRT, likelihood ratio testing. p.-h., post-hoc tests. *A. f.*, *A. franciscana*. *A. p.*, *A.* *parthenogenetica*. sp., species. \*, interactions between the factors were included. NA, not applicable.

Analyses were run in R version 3.4.2 (R Core Team 2014) using the packages lme4 (linear mixed models, Bates et al. 2015), survival (survival analyses, Therneau 2014), pscl (hurdle models, Zeileis et al. 2008), and multcomp (fuction “glht” for post-hoc testing, Hothorn et al. 2008).

To estimate the infectiousness of a single spore, we used the results of the transmission assay. We assumed that the establishment of microsporidian infections follows an independent-action model with birth-death processes. This model assumes that a parasite population grows in the host until it reaches an infective threshold, at which point the infection is considered to be established (Schmid-Hempel 2011 pp. 225–6). In our assay, we considered that an infection was established when we could detect it; in other words, the infective threshold corresponded to the threshold for PCR detection (estimated at ∼1000 spores inside the host’s body, unpublished data). In these models, the probability per spore to start an infection, *p*, is equal to 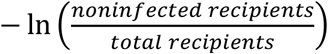 /*D* where *D* is the spore dose (Schmid-Hempel 2011 pp. 225–6). In our transmission assay, *D* can be approximated by the number of spores in the fecal sample taken from the donor at the start of the assay (= spore count transformed to spores/mL, or * 700), divided by 5*8=40 (fecal samples accumulated over a 5-day period but we only exposed recipients for one day; the inoculum was shared amongst 8 recipients). We calculated a value of *p* for every replicate in the transmission assay.

For each infection we used two measures of spore production as proxies for parasite fitness. First, we calculated a proxy for the ‘lifetime transmission success’: we summed the number of spores in the fecal samples taken on days 15, 30, 45 and 60 for each infection, then corrected this cumulative spore count by *p*, the average infectiousness of a single spore in a given host-parasite combination (as calculated above). Second, we calculated an asymptotic growth rate by computing the dominant eigenvalue of a standard Leslie matrix,

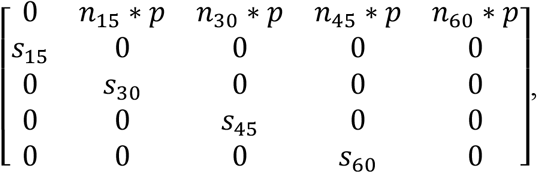

where *n_i_* is the number of spores in the fecal sample on day *i*, *p* is the average infectiousness of a single spore in that host-parasite combination (as calculated above), and *s_i_* describes whether the host survived until day *i* (1) or not (0). While the lifetime transmission success is a measure of the basic reproduction number R_0_, which describes parasite fitness under stable endemic conditions, the asymptotic growth rate is a measure of the net population growth rate, which describes fitness under epidemic conditions (Frank 1996, Hethcote 2000); we included both measures because either situation can occur in the field.

#### Statistical analyses: infection vs. resistance

In most of the experimental host-parasite combinations, a subset of exposed hosts did not become (detectably) infected. Hereafter, we refer to these individuals as resistant, because we found *a posteriori* differences in the proportion of such individuals across host-parasite combinations, and in their life history traits compared to infected individuals and controls. As above, the analyses of these two aspects excluded all individuals who died before infection status could be definitively determined, i.e. those that died before day 15 of the experiment.

We analyzed the distribution of resistance across host-parasite combinations. Within each host species, we used χ^2^ tests to compare the numbers of resistant and infected hosts after exposure to *A. rigaudi* and *E. artemiae*. We also used χ^2^ tests to test for an effect of sex on the probability of resistance to each parasite in *A. franciscana*, and for an effect of spore dose on the probability of resistance to *E. artemiae* in *A. parthenogenetica*.

There was substantial variation in infection outcome for the combinations *A. franciscana*-*A. rigaudi*, and *A. parthenogenetica*-*E. artemiae* (low dose) (see Results). Because costs of resistance are a common aspect of host-parasite interactions (Schmid-Hempel 2003), we investigated whether resistance was related to host fitness in these combinations. To do this, we repeated the survival and reproduction analyses described above, with an added *Resistant-Infected-Control* factor. We added or excluded this factor and its interactions with the other fixed effects, then compared all models using the corrected AIC. In this way, we investigated whether the outcome of infection explained a significant part of the variation in host traits after the experimentally manipulated factors were taken into account. If the *Resistant-Infected-Control* factor was maintained in the best models, we used contrast manipulation and AICc-based model comparison to detect how the three host categories (*Resistant*, *Infected*, *Control*) differed.

## Results

### Experiment 1: Infectivity

Both *A. parthenogenetica* and *A. franciscana* were more susceptible to infection with *A. rigaudi* than *E. artemiae* (χ^2^(3)≥20.9, *p*<0.001 for both; Fig. 1). For *A. franciscana*, the slopes and inflection points of the two curves were not significantly different, but the upper limit was significantly higher for *A. rigaudi* than for *E. artemiae* (*t*=2.1, *p*=0.03). In *A. parthenogenetica*, the infectivity of the two parasites was markedly different: successful infections with *E. artemiae* required such a high spore dose that the inflection point and upper limit of its curve could not be computed; its slope was not significantly different to that of *A. rigaudi*. Mortality was not dose-dependent in any of the host-microsporidian combinations, so we can be confident that it did not skew results (Supp. Table 1).

**Figure 1.**
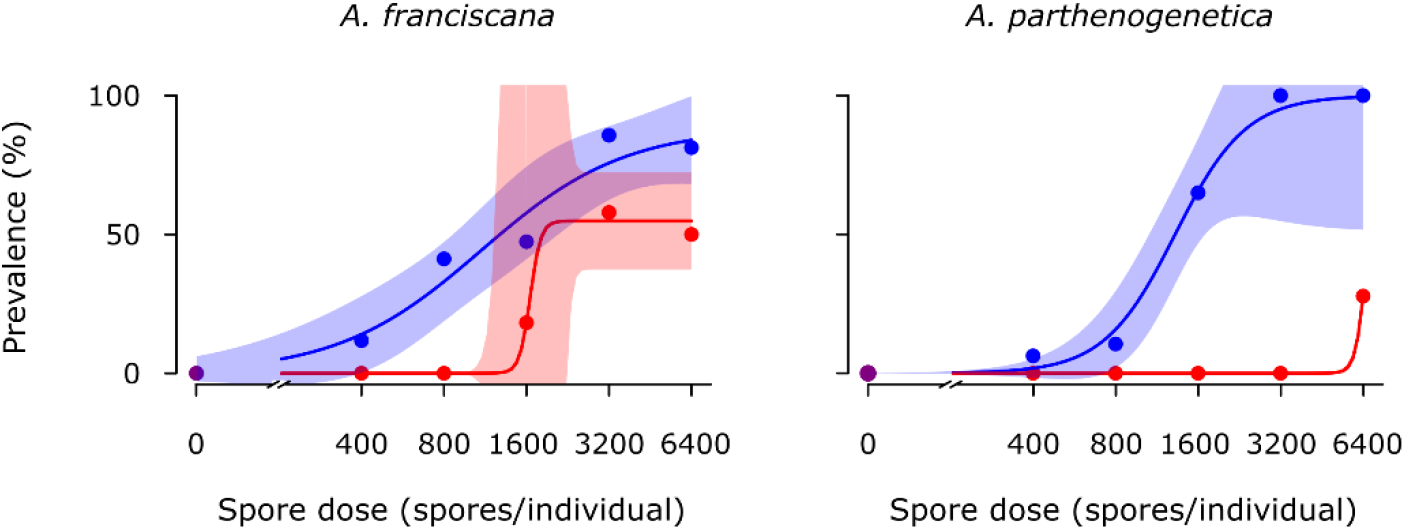
Infectivity of *A. rigaudi* (blue) and *E. artemiae* (red) in *A. franciscana* (left) and *A. parthenogenetica* (right). Points indicate the prevalence (% infected) at each dose; lines are the best fits and the shaded areas represent the 95% CIs. Because the inflection point of *E. artemiae* in *A. franciscana* was poorly resolved, uncertainty was high here. It was not possible to calculate a confidence interval for *E. artemiae* in *A. parthenogenetica* due to low resolution.

### Experiment 2: Virulence and transmission

Among host individuals that survived until we could be certain of their infection status (i.e. that survived until at least day 15), infection rates were high (Table 3). As expected, many fewer infections were detected among individuals that died before day 15. In general, infection rates in Experiment 2 were considerably higher than those in Experiment 1; this was most likely because the longer incubation time allowed slow-growing infections to become detectable.

**Table 3.** Detection of infection before and after the detection threshold. The table shows the detected infection rates in individuals that survived until we could be certain of their infection status (i.e. that died after the detection threshold on day 15) vs. individuals that died before this threshold day.

| Host-parasite combination |  | Infection rate after vs. before the detection threshold |  |
| --- | --- | --- | --- |
| ***A. franciscana*** |  |  |  |
| Exposure to *A. rigaudi* |  | 86 % | 50 % |
| Exposure to *E. artemiae* |  | 96 % | 13 % |
| ***A. parthenogenetica*** |  |  |  |
| Exposure to *A. rigaudi* |  | 100 % | 15 % |
| Exposure to *E. artemiae* | – low spore dose | 64 % | 0 % |
|  | – high spore dose | 86 % | 20 % |

#### Virulence of infections

We analyzed the species-level results of Experiment 2 in two parts. First, we analyzed the virulence of parasite infections, expressed as effects on host survival, growth, and reproduction. These results are summarized in Fig. 2 and the significance of tested effects is listed in Table 4; we discuss the effects of infection in more detail below. Here, we report only the analyses for infected vs. control individuals, which excluded all individuals that died before day 15. When we compared exposed vs. control individuals that died before the cut-off day the results were not qualitatively different.

**Figure 2.**
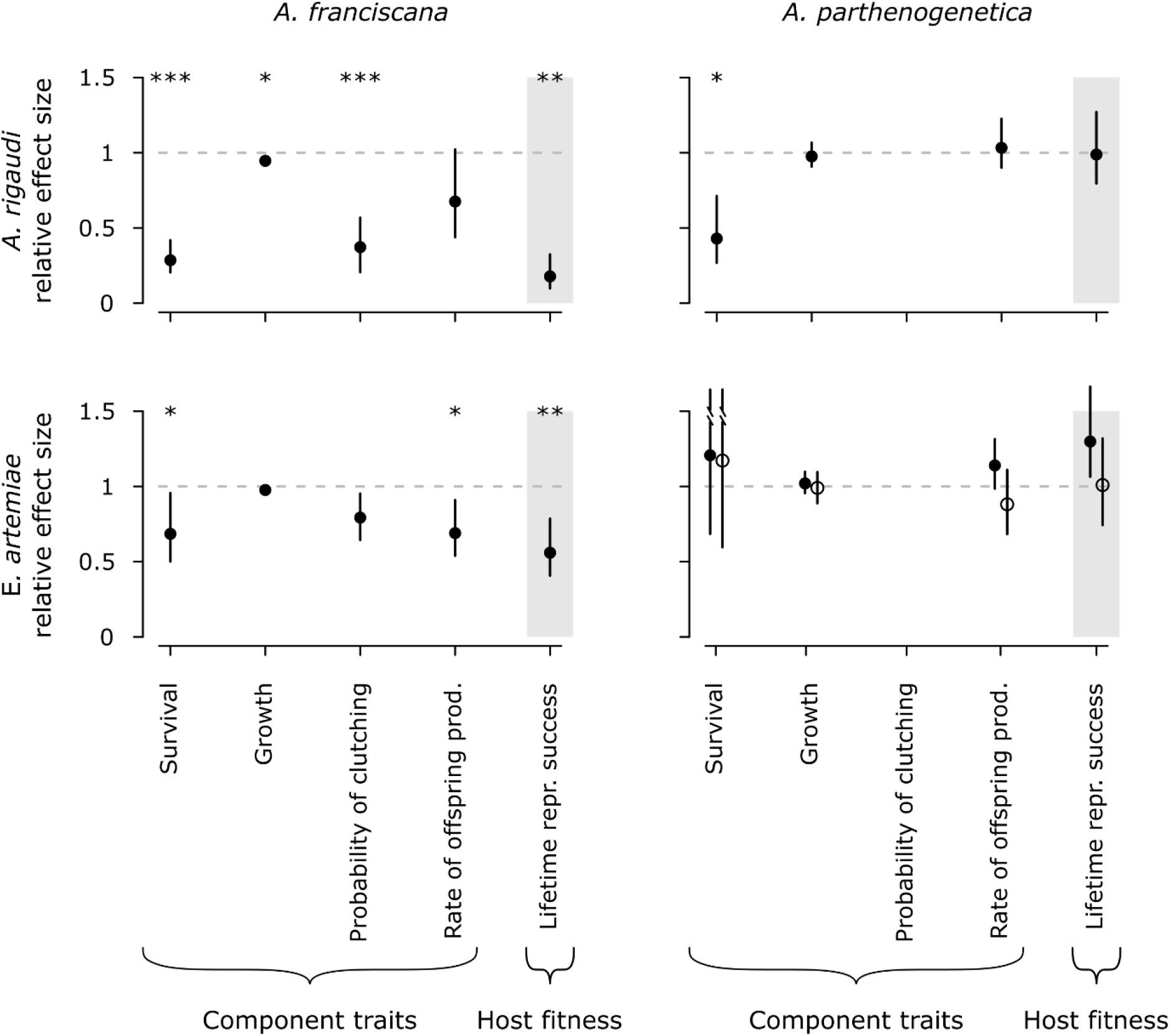
Host fitness (≈ parasite virulence) in the four host-parasite combinations. All factors are shown as fitted effects relative to controls: survival is an acceleration factor (the ratio of expected time-until-death); the probability of reproduction is a relative risk; growth, rate of offspring production, and LRS are ratios. Bars represent the 95% profile likelihood CIs (survival) or bootstrapped CIs (all others). *A. parthenogenetica* infected after exposure to 10000 *E. artemiae* spores are indicated with open circles. Asterisks indicate significant differences from controls (represented by the dotted gray line). The plotted survival effect for *A. parthenogenetica* excludes the aberrant group (see Results). All reproductive and fitness traits were obtained for females only. The probability of reproduction is not shown for *A. parthenogenetica* because it could not be analyzed. Weighing the contributions of nauplii and cysts to the rate of offspring production and LRS generated qualitatively equivalent results; the results shown here are for equal weights.

**Table 4.** Significance of tested effects for the virulence of infections. Analyses were run separately for *A. franciscana* and *A. parthenogenetica*. See text for post-hoc analyses of Treatment.

| Tested variable | Fixed-effect terms | test statistic, *p* | Effect |
| --- | --- | --- | --- |
| **Virulence of infections: *A. franciscana*** |  |  |  |
| **Survival** | Treatment | $\chi^2(2) = 48.2, p < 0.0001$ | ↓ when infected |
| | Sex | $\chi^2(1) = 33.2, p < 0.0001$ | ↑ for males |
| | Size class | $\chi^2(1) = 4.3, p = 0.04$ | ↑ for larger individuals |
|  | interactions | all non-significant |  |
| **Growth between days 1 & 30** | Treatment | $\chi^2(2) = 9.7, p < 0.01$ | |
| | Sex | $\chi^2(1) = 133.5, p < 0.0001$ | ↓ for males |
| | Size class | $\chi^2(1) = 95.0, p < 0.0001$ | ↓ for larger individuals |
|  | interactions | all non-significant |  |
| **Reproduction** |  |  |  |
| Time until sexual maturity | Treatment | $\chi^2(2) = 22.5, p < 0.0001$ | ↑ when infected |
| | Size class | $\chi^2(1) = 0.1, p = 0.83$ | |
| | interaction | $\chi^2(2) = 1.3, p = 0.54$ | |
| Probability of producing a clutch | Treatment | $\chi^2(2) = 31.3, p < 0.0001$ | ↓ when infected |
| | Size class | $\chi^2(1) = 0.5, p = 0.50$ | |
| | interaction | $\chi^2(2) = 0.8, p = 0.69$ | |
| Rate of offspring production | Treatment | $\chi^2(2) \geq 7.9, p \leq 0.02$ <sup>†</sup> | ↓ when infected |
| | Size class | $\chi^2(1) \leq 0.5, p \geq 0.50$ <sup>†</sup> | |
| | interaction | $\chi^2(2) \leq 1.8, p \geq 0.41$ <sup>†</sup> | |
| Timing of offspring production | Treatm. : % Repr. period | $\chi^2(4) \leq 3.7, p \geq 0.45$ <sup>†</sup> | |
| Type of offspring produced | Treatment | $\chi^2(2) = 16.8, p < 0.001$ | more nauplii when infected |
| | Size class | $\chi^2(1) = 0.1, p = 0.73$ | |
| | interaction | $\chi^2(2) = 0.7, p = 0.71$ | |
| **Fitness (LRS)** | Treatment | $\chi^2(4) \geq 46.6, p < 0.0001$ <sup>†</sup> | ↓ when infected |
| | Size class | $\chi^2(2) \leq 1.5, p \geq 0.48$ <sup>†</sup> | |
| | Interaction | $\chi^2(4) \leq 1.5, p \geq 0.82$ <sup>†</sup> | |
| **Virulence of infections: *A. parthenogenetica*** |  |  |  |
| **Survival** | Treatment | $\chi^2(3) = 19.7, p < 0.001$ | ↓ when infected |
| | Size class | $\chi^2(2) = 11.5, p < 0.01$ | ↑ for larger individuals |
|  | interaction | see text |  |
| **Growth between days 1 & 30** | Treatment | $\chi^2(3) = 1.3, p = 0.73$ | |
| | Size class | $\chi^2(2) = 35.8, p < 0.0001$ | ↓ for larger individuals |
| | interaction | $\chi^2(6) = 13.1, p = 0.04$ | see text |
| **Reproduction** |  |  |  |
| Rate of offspring production | Treatment | $\chi^2(3) \leq 5.8, p > 0.12$ <sup>†</sup> | |
| | Size class | $\chi^2(2) \geq 8.8, p = 0.01$ <sup>†</sup> | ↑ for larger individuals |
| | interaction | $\chi^2(6) \leq 10.4, p \geq 0.11$ <sup>†</sup> | |
| Timing of offspring production | Treatm. : % Repr. period | $\chi^2(4) \geq 10.4, p < 0.11$ <sup>†</sup> | earlier when infected |
| Type of offspring produced | Treatment | $\chi^2(2) = 1.4, p = 0.71$ | |
| | Size class | $\chi^2(2) = 0.1, p = 0.96$ | |
| | interaction | $\chi^2(6) = 9.1, p = 0.17$ | |
| **Fitness (LRS)** | Treatment | $\chi^2(6) \leq 5.6, p \geq 0.13$ <sup>†</sup> | |
| | Size class | $\chi^2(4) \geq 8.2, p < 0.09$ <sup>†</sup> | ↓ for largest individuals |
| | interaction | $\chi^2(12) \leq 13.8, p \geq 0.31$ <sup>†</sup> | |

<sup>†</sup>Depending on the weight of nauplii vs. cysts.

In most host-parasite combinations, survival was reduced (Fig. 2, Table 4). For *A. franciscana*, a lognormal survival model best fit the data (ΔAICc≥4.2). Infection significantly reduced survival; post-hoc testing revealed that this effect was highly significant for *A. rigaudi* and marginally significant for *E. artemiae* (*t*=-6.7 and −2.2, *p*<0.0001 and *p*=0.05, respectively). For *A. parthenogenetica*, a log-logistic survival model best fit the data (ΔAICc≥0.9). Survival was affected by infection, size class, and their interaction, but this complicated interaction effect was due to the aberrant survival curves of one group of individuals (Batch 34 ± 2 days old, Size class 7.5 mm), which had high death rates for controls and low death rates for infected hosts. When this group was removed, the interaction effect became non-significant. In general therefore, survival of *A. parthenogenetica* was reduced by infection with a parasite; post-hoc testing revealed that individuals infected with *A. rigaudi* had significantly lower survival (*t*=-3.3, *p*<0.01), while individuals infected with *E. artemiae* did not (*t*=0.7 and 0.5, *p*=0.86 and 0.94 for low and high spore dose, respectively).

More than 90% of host growth occurred between days 1 and 30 (Supp. Table 2), so only this period was analyzed (Fig. 2, Table 4). For *A. franciscana*, infection significantly reduced growth; this effect was driven by *A. rigaudi* (post-hoc *z*=-3.1, *p*<0.01) and non-significant for *E. artemiae* (post-hoc *z*=-1.5, *p*=0.23). *A. parthenogenetica* growth was affected by infection interacting with size class, with the smallest size class growing slightly less when infected with a high dose of *E. artemiae*, and the middle size class growing slightly more when infected with either dose of *E. artemiae*.

Parasite infection affected the reproduction of *A. franciscana* females in various ways (Fig. 2, Table 4). The time until maturity, which was best described by a lognormal distribution (ΔAICc≥4.7), was significantly delayed by infection with either parasite species (post-hoc for *A. rigaudi t*=4.6, *p*<0.0001; post-hoc for *E. artemiae t*=2.5, *p*=0.02). The probability that *A. franciscana* females produced a clutch was also significantly lower when they were infected; this effect was driven by *A. rigaudi* (post-hoc *z*=-5.3, *p*<0.0001) and was marginally non-significant for *E. artemiae* (post-hoc *z*=-2.1, *p*=0.06). For *A. franciscana* females that did reproduce, infection with *A. rigaudi* increased the proportion of nauplii clutches (post-hoc for *A. rigaudi z*=3.9, *p*<0.001; post-hoc for *E. artemiae z*=0.8, *p*=0.68). The rate of offspring production was significantly reduced by infection with *E. artemiae* for all weights of nauplii vs. cysts (post-hoc for *E. artemiae z*≤-2.8, *p*≤0.01), but was only significantly reduced by infection with *A. rigaudi* when cysts were weighted twice as much as nauplii (*z*=-2.3, *p*=0.04). Finally, the timing of offspring production was independent of infection status.

In contrast, parasite infection had little effect on the reproduction of *A. parthenogenetica* females (Fig. 2, Table 4). The effects of infection on the time until sexual maturity or the probability of producing a clutch could not be tested, because almost all *A. parthenogenetica* females started reproducing immediately. For reproducing females, neither the proportion of live clutches, nor the rate of offspring production were affected by infection with either parasite. However, infection with *A. rigaudi* did lead to a significant shift towards earlier reproduction (Fig. 3; significant effect of treatment when cysts were weighted equally or doubly compared to nauplii; post-hoc for *A. rigaudi z*≥2.6, *p*≤0.03 for all weights of nauplii vs. cysts; post-hoc for *E. artemiae z*≤1.1, *p*≥0.58 for all weights of nauplii vs. cysts).

**Figure 3.**
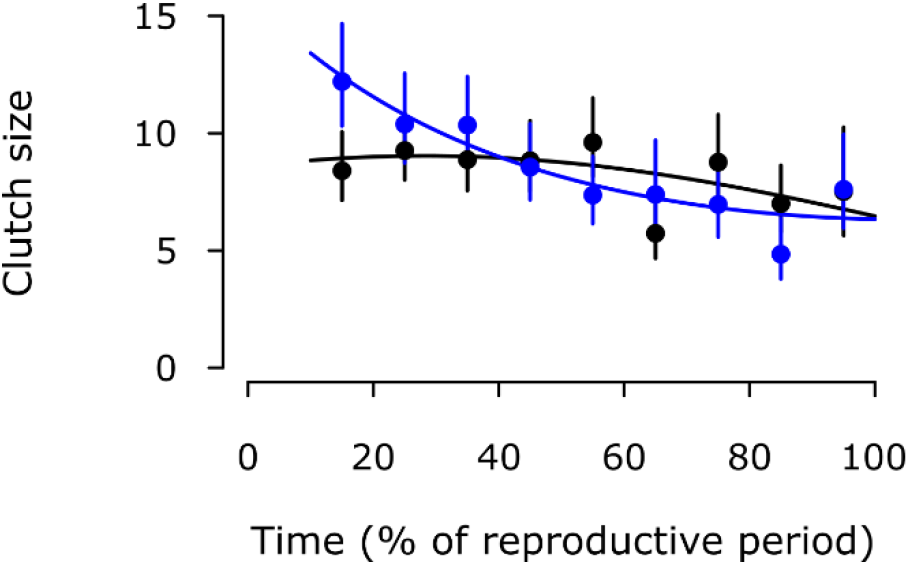
Timing of reproduction in *A. parthenogenetica* controls (black) and infected with *A. rigaudi* (blue). Lines represent the prediction of the best model, points and vertical bars give the observed means and their 95% CIs, calculated over intervals of 10%. Weighing the contributions of nauplii and cysts to the total number of offspring generated qualitatively similar results; the results shown here are for equal weights.

The fitness of female hosts – estimated by the lifetime reproductive success (LRS), i.e. the total number of offspring produced – was significantly reduced by infection with either parasite for *A. franciscana* (post-hoc for *A. rigaudi t*≤-7.3, *p*≤0.0001 for all weights of nauplii vs. cysts; post-hoc for *E. artemiae t*≤-3.9, *p*≤0.001 for all weights of nauplii vs. cysts), but not for *A. parthenogenetica* (Fig. 2, Table 4).

#### Transmission and fitness of infections

Second, we studied the effects of the host species on the parasite’s transmission and fitness (summarized in Fig. 4). These analyses were combined for all host-parasite combinations, but *A. parthenogenetica* that were exposed to 10000 *E. artemiae* spores were analyzed separately unless otherwise specified.

**Figure 4.**
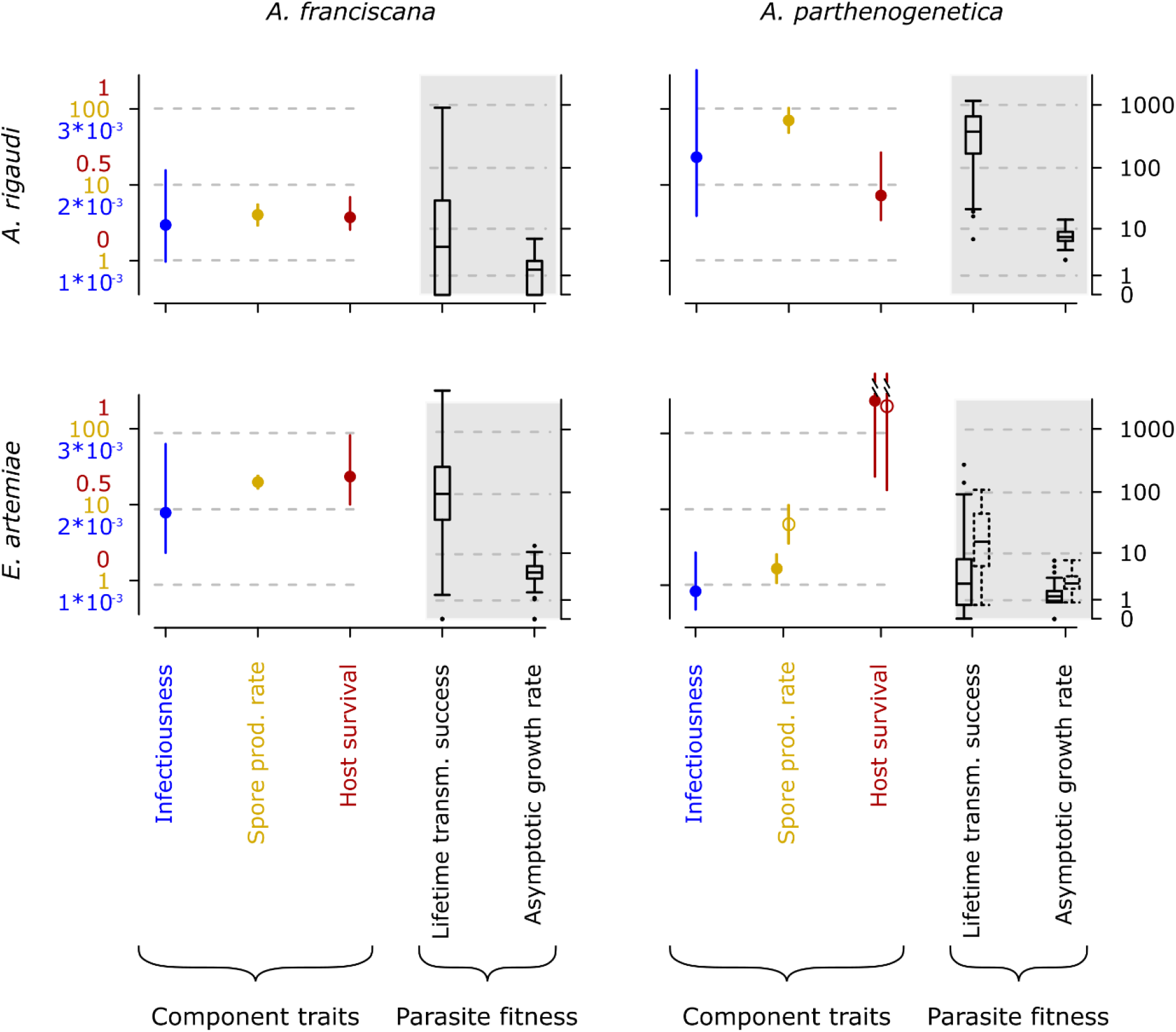
Parasite fitness in the four host-parasite combinations. The component traits infectiousness (probability of infection by a single spore), rate of spore production (# counted spores/5 days, *ln* scale), and host survival (which determines infection duration, copied from Fig. 2) are shown as fitted means with 95% profile likelihood CIs. The fitness measures lifetime transmission success (*ln + 1* scale) and asymptotic growth rate (*ln + 1* scale) are shown as Tukey box plots. *A. parthenogenetica* infected after exposure to 10000 *E. artemiae* spores are indicated with open circles and dotted box plots. Note that spore production, host survival, and parasite fitness were analyzed for infected hosts only.

The infectiousness of a single spore (the probability that it started a detectable infection, as calculated using the transmission data) corresponded with our expectations based on Experiment 1 (Fig. 4). Host-parasite combination had a significant effect on infectiousness (χ^2^(1)=16.7, *p*<0.0001). *A. rigaudi* tended to be more infectious to *A. parthenogenetica* than to *A. franciscana* (post-hoc *z*=2.3, *p*=0.10); *E. artemiae* was significantly more infectious to *A. franciscana* than to *A. parthenogenetica* (post-hoc *z*=3.6, *p*<0.01).

The rates of spore production were significantly different in all host-parasite combinations (overall χ^2^(1)=205.9, *p*<0.0001; all post-hoc pairwise comparisons *z*≥3.2, p<0.01; Fig. 4); they were highest in the combinations *A. parthenogenetica-A. rigaudi* and *A. franciscana-E. artemiae*. For *A. parthenogenetica* infected with *E. artemiae*, the rate of spore production was notably higher when the initial inoculum was larger (χ^2^(1)=10.6, *p*=0.001).

As expected, host-to-host transmission success increased with the rate of spore production in all host-parasite combinations (Spearman’s *ρ* between 0.57 and 0.69, *p*<0.0001; Supp. Fig. 1). Therefore, we were able to use the lifetime transmission success and asymptotic growth rate as indicators of parasite fitness. The two measures were tightly correlated (Supp. Fig. 2) and both differed across host-parasite combinations (χ^2^(4)=189.9 and 245.0, respectively, *p*<0.0001; Fig. 4). The fitness of *A. rigaudi* infections was highest in *A. parthenogenetica*; that of *E. artemiae* infections was highest in *A. franciscana.* All pairs of host-parasite combinations were significantly different, except the low-performers *A. parthenogenetica-E. artemiae* and *A. franciscana-A. rigaudi* (post-hoc *p*<0.001 vs. *p*=0.46 and *p*≤0.04 vs. *p*=0.69, respectively). This was true for both spore doses of *A. parthenogenetica-E. artemiae* regarding the lifetime transmission success, but only for the low spore dose for the asymptotic growth rate.

#### Infection vs. resistance

Among the individuals that survived until we could be certain of their infection status, the rate of resistance varied between 0 and 36% in the different host-parasite combinations (Table 5). For *A. franciscana*, significantly more individuals resisted infection with *A. rigaudi* than with *E. artemiae* (χ^2^(1)=10.4, *p*<0.01), and this effect was independent of sex (χ^2^(1)=0.3, *p*=0.58). Similarly, significantly more *A. parthenogenetica* resisted infection with *E. artemiae* (χ^2^(1)=20. 6, *p*<0.0001), with a marginally non-significant difference between the two spore doses (χ^2^(1)=3.8, *p*=0.052). There was substantial variation in infection outcome for the combinations *A. franciscana*-*A. rigaudi*, and *A. parthenogenetica*-*E. artemiae* (low dose), so we continued our analyses with these combinations.

**Table 5.** Number of exposed individuals that were infected or resistant to *A. rigaudi* and *E. artemiae*. These counts excluded all individuals who died before infection status could be definitively determined, i.e. those that died before day 15 of the experiment.

| Host-parasite combination | Resistant | Infected | % Resistant |
| --- | --- | --- | --- |
| ***A. franciscana* males** |  |  |  |
| Exposure to *A. rigaudi* | 8 | 59 | 12% |
| Exposure to *E. artemiae* | 5 | 106 | 5% |
| ***A. franciscana* females** |  |  |  |
| Exposure to *A. rigaudi* | 12 | 60 | 17% |
| Exposure to *E. artemiae* | 5 | 114 | 4% |
| ***A. parthenogenetica*** |  |  |  |
| Exposure to *A. rigaudi* | 0 | 62 | 0% |
| Exposure to *E. artemiae* – low spore dose | 27 | 49 | 36% |
| – high spore dose | 4 | 25 | 14% |

For both *A. franciscana* exposed to *A. rigaudi* and *A. parthenogenetica* exposed to a low spore dose of *E. artemiae*, resistant individuals died more quickly than infected individuals (Fig. 5; ΔAICc respectively>4.4 and=1.6, Supp. Table 3). For *A. franciscana*, resistant males had a higher mortality than resistant females (Fig. 5; ΔAICc>1.7, Supp. Table 3).

**Figure 5.**
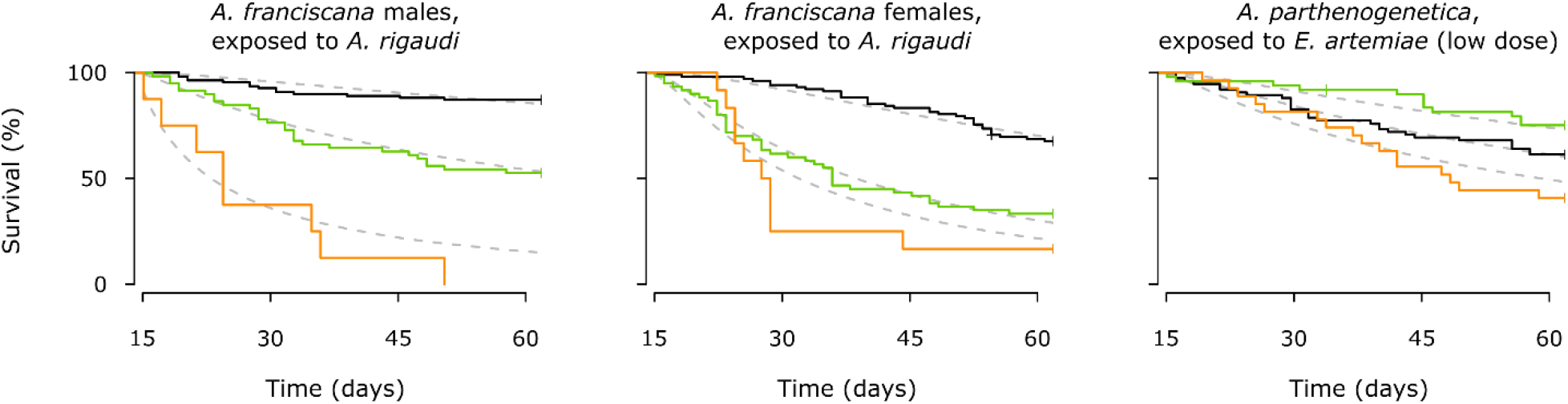
Survival curves for resistant (orange), infected (green), and control (black) individuals. Note that these curves start at day 15, i.e. when infection status could be fully ascertained. The curves shown here are averaged across size class and origin for *A. franciscana* and across size classes in *A. parthenogenetica*. Model estimates for each curve are plotted in gray.

Finally, there was little support for an effect of resistance on reproduction in females of either host species (Supp. Table 4). *A. franciscana* females that resisted infection with *A. rigaudi* behaved similarly to females that became infected (strong effects of *Resistant-Infected-Control*, but no or weak support for a difference between resistant and infected females). The reproductive behavior of *A. parthenogenetica* females that resisted infection with a low dose of *E. artemiae* was similar to that of infected and control females.

## Discussion

The degree of host specialization is a key property of any multi-host parasite. Host specialization, when considered as a difference in fitness, arises from a series of life history traits including the ability to infect, the rate of transmission, and the virulence. We quantified these traits for two microsporidian gut parasites (*A. rigaudi* and *E. artemiae*) infecting two brine shrimp hosts (*A. franciscana* and *A. parthenogenetica*), by tracking the life history of both hosts and parasites after experimental infection. A brief synopsis of the results is shown in Table 6.

**Table 6.** Qualitative synopsis of results.

| Host species | Parasite species |  |
| --- | --- | --- |
|  | *A. rigaudi* | *E. artemiae* |
| ***A. franciscana*** | moderately infectious<br>low spore production<br>highly virulent<br>⇒ **low parasite fitness,**<br>**mismatched host & parasite** | highly infectious<br>high spore production<br>moderately virulent<br>⇒ **high parasite fitness,**<br>**matched host & parasite** |
| ***A. parthenogenetica*** | highly infectious<br>high spore production<br>moderately virulent (survival only)<br>⇒ **high parasite fitness,**<br>**matched host & parasite** | poorly infectious<br>low spore production<br>avirulent<br>⇒ **low parasite fitness,**<br>**mismatched host & parasite** |

Overall, each of the parasites was partially specialized: *A. rigaudi* was very successful in *A. parthenogenetica*, while *E. artemiae* performed best in *A. franciscana*. Below, we discuss how the individual life-history traits combine to shape the degree of specialization, and the ensuing effects of specialization on the hosts. We refer to the host-parasite combinations where parasites reached high fitness as the ‘matched’ combinations (Table 6). The reversed combinations also produced viable transmission stages, but at much lower rates; we will call these the ‘mismatched’ combinations.

### Partial specialization via a mix of specialist and generalist traits

Specialization is often presented as a dichotomy: specialists, whose fitness is high or null for different hosts, versus generalists, who generally have intermediate fitness on several hosts (Poulin 2007 chap. 3, Schmid-Hempel 2011 chap. 7, Leggett et al. 2013). *A. rigaudi* and *E. artemiae* fall into a gray zone between these categories, being neither absolute specialists – they can exploit both hosts –, nor absolute generalists – their fitness is much higher in the matched hosts.

When broken down into its component traits, the origin of this partial specialization becomes clear. Parasites should be as infective as possible to hosts to which they are adapted, and indeed both *A. rigaudi* and *E. artemiae* are highly infectious to their matched hosts. Similarly, we expect strong transmission to be advantageous, and accordingly we find that both parasites have high rates of spore production in their matched hosts. The expectations for virulence are not as clear-cut. A ‘Darwinian devil’ parasite would be avirulent while maintaining high transmission rates, but it is generally considered that these two factors are correlated (Alizon et al. 2009). Virulence must therefore be judged in relation to transmission; for example, high virulence can be adaptive if coupled with high rates of transmission, or maladaptive if not. When considered in this way, *A. rigaudi* and *E. artemiae*’s virulence are also coherent with their overall specialization. *A. rigaudi* causes high survival virulence – and thus short infection durations – in both hosts, but this is advantageously coupled with high rates of spore production in its matched host *A. parthenogenetica*, and disadvantageously coupled with low rates of spore production in its mismatched host. *E. artemiae* is avirulent in its mismatched host, which at first glance appears ideal. However, when spore production is taken into account, it becomes clear that this avirulence in *A. parthenogenetica* is coupled with very low rates of transmission, whereas the rate of spore production is high in *A. franciscana*.

Despite this, were we to consider the component traits individually, they would not all lead us to conclude that the two parasites are partially specialized. The pattern of spore production in the matched vs. mismatched combinations best reflects the overall degree of specialization. Infectivity, on its own, might lead us to conclude that *E. artemiae* is a specialist while *A. rigaudi* is more generalist (also discussed in Lievens et al. subm.). Virulence is difficult to interpret outside the context of spore production, as discussed above, making it a particularly poor proxy for overall specialization. Integrating across all of these life history traits is therefore necessary to properly understand the nature of this host-parasite system, and will probably have important implications for the evolution of virulence (Alizon and Michalakis 2015) and infection success (Hall et al. 2017).

### Mismatched parasites have different kinds of suboptimal virulence

Several theoretical predictions have been made for the evolution of virulence in multi-host parasites that are specialized on one host and spill over into another (source-sink dynamics), all of which agree that virulence should depend exclusively on the optimum in the specialized host (Regoes et al. 2000, Woolhouse et al. 2001, Dobson 2004, Gandon 2004). Predictions of virulence in the non-specialized host, however, vary. Regoes et al. considered virulence to be coupled to exploitation, which trades off between hosts; their prediction is that the parasite will be avirulent in the spillover host. Gandon also considered virulence to be coupled to exploitation, but in his model the level of exploitation is correlated between hosts. In this case, the parasite can be maladaptively avirulent or hypervirulent in the spillover host, depending on the relative resistances of the hosts. Finally, Woolhouse et al. pointed out that virulence can become decoupled from parasite exploitation in spillover hosts, for example through harmful immune responses (Graham et al. 2005), leading to maladaptively high virulence (see also Leggett et al. 2013). Empirically, virulence patterns across multiple hosts have only rarely been studied in natural systems (Rigaud et al. 2010), so it is difficult to determine which of these possibilities may be more common.

In the mismatched hosts of our *Artemia*-microsporidian system, two different virulence patterns are apparent. First, in the combination *A. franciscana-A. rigaudi*, the parasite is very virulent on a host in which it can barely reproduce. Its virulence in the non-specialized host is thus decoupled from exploitation and maladaptive, matching Woolhouse et al. (2001)’s prediction for unconstrainedly high virulence. The situation of *A. rigaudi* strongly resembles that of the generalist microsporidian parasite *Nosema bombi*, which infects bumble bees. Two of *N. bombi*’s most important hosts are *Bombus terrestris*, in whom it is so virulent that it cripples its own year-to-year transmission, and *Bombus lucorum*, in whom its virulence is moderate enough to allow transmission (Rutrecht and Brown 2009). A number of zoonotic human diseases also fit this pattern (Woolhouse et al. 2001, cf. Auld et al. 2017). In contrast, in the mismatched combination *A. parthenogenetica-E. artemiae*, the parasite is avirulent. *E. artemiae* could therefore correspond to the situations described by Regoes et al. (2000) and Gandon (2004), in which a non-specialized host is under-exploited and suffers no virulence. Indeed, *A. parthenogenetica* is also less susceptible to *E. artemiae*, giving some support to Gandon’s scenario of differently resistant hosts. A similar case could be made for the nematode *Howardula aoronymphium* (Jaenike 1996, Jaenike and Dombeck 1998, Perlman and Jaenike 2003) and for the Drosophila C virus (Longdon et al. 2015), which exhibit a range of exploitation and correlated virulence across host species.

Overall, our results provide support for the varied possible theoretical predictions of virulence evolution in multi-host parasites: in one case, we appear to be dealing with decoupled, ‘runaway’ virulence, while in the second the differences in virulence may be driven by levels of host resistance. These contrasting findings show that the different theoretical outcomes can even be found among host-parasite pairs that are ecologically extremely similar and phylogenetically close.

### Mismatched hosts incur high costs of resistance

In the matched host-parasite combinations, uninfected individuals were rare or nonexistent (Table 5), and suffered no detectable survival cost (data not shown). It is possible that an extremely high mortality rate of resistant individuals caused them to die before we could reliably detect infection, leading us to underestimate both the frequency and the cost of resistance. However, survival rates for the matched combinations were universally high in the infectivity experiment, which lasted one week. Any mortality conferred by resistance would therefore have to be incurred precisely in the second week of infection, which is unlikely. It is more probable that the high rates of infection reflect selection on the parasite to evade or overcome resistance in its matched host (Hasu et al. 2009).

In the mismatched host-parasite combinations, however, up to one third of the exposed hosts were uninfected, and the life histories of these individuals differed clearly from those of control or infected hosts (Table 5, Fig. 5). This suggests that their lack of infection was the result of an active resistance mechanism. Because the parasite was absent, the effects of deploying resistance must have been induced by the host itself, as a consequence of its immune reaction upon exposure (immunopathology, Schmid-Hempel 2003, Graham et al. 2005).

This resistance was extremely costly: resistant individuals died much more rapidly than control and infected hosts (Fig. 5). Since there was no detectable compensation through increased fecundity, we must conclude that resistance in these cases is maladaptive. This is intriguing, because *A. franciscana* and *A. parthenogenetica* are regularly exposed to their mismatched parasites in the field (Lievens et al. subm.). Host resistance has been shown to evolve quickly in a similar host-parasite system (*Daphnia magna-Octosporea bayeri*, Zbinden et al. 2008), so we would not expect maladaptive resistance responses to persist in the host populations. An explanation may be that source-sink dynamics acting in the parasite populations prevent them from evolving to reduce their impact on the mismatched hosts. In turn, selection on the host to reduce its response to the mismatched parasite could perhaps be countered by other factors, such as the need to maintain its overall immune capacity (Graham et al. 2005). Similarly disproportionate costs of resistance, with uninfected hosts dying more rapidly than even infected hosts, have been found in e.g. *Daphnia* resisting the bacterium *Pasteuria* (Little and Killick 2007, though see Labbé et al. 2010), and naïve isopods resisting infection with a helminth (Hasu et al. 2009).

### Infection with A. rigaudi causes shifts in reproductive strategy

*A. parthenogenetica* females infected with their matched parasite *A. rigaudi* died more quickly than controls and did not produce offspring at a higher overall rate, yet did not suffer from reduced lifetime reproductive success. They managed this by shifting towards earlier reproduction to alleviate the survival virulence, a plastic behavior known as fecundity compensation (cf. Minchella and Loverde 1981, Agnew et al. 2000, Chadwick and Little 2005) (Fig. 3). Females accomplished this shift in reproductive effort by increasing the size, rather than the frequency, of early clutches (frequency data not shown). This is a new finding for *Artemia*, which opens a number of interesting new avenues of research. For example, Mediterranean *A. parthenogenetica* are also heavily infected with the castrating cestode *Flamingolepis liguloides* (Amat et al. 1991). If fecundity compensation is also possible in the face of *F. liguloides* infections, this would drastically change our understanding of the overall impact of this parasite. Alternatively, a recent study has shown that the fecundity compensation response of snails can be suppressed when they undergo additional stress (Gleichsner et al. 2016). Studying the relationship between common environmental stressors of *A. parthenogenetica*, such as salinity, and their ability to shift their reproductive effort may help us understand the evolutionary relevance of such mechanisms.

*A. franciscana* females did not have a similar fecundity compensation response when infected with either parasite. However, infections of *A. franciscana* with *A. rigaudi* were associated with an interesting change in reproductive strategy. Infected females were less likely to produce a clutch, but those that did reproduce were more likely to produce clutches of live young. Considering that *Artemia* generally produce cysts when stressed (Clegg and Trotman 2002), this result seems counterintuitive. Perhaps *A. rigaudi* interferes with the cyst production mechanism, either collaterally or as a manipulation to increase the availability of susceptible hosts. Another possibility is that a shift towards live born offspring is advantageous for the host. If infected mothers can produce offspring that are protected against the parasite, for example via transgenerational immune priming (which Artemia can do, Norouzitallab et al. 2015), those offspring should have a competitive advantage when encountering the parasite. If this protection is costly, it may be more worthwhile to produce protected nauplii than protected cysts: protected nauplii will certainly be born into a parasite-infested environment, while the hatching environment of protected cysts is unknown.

### Comparison with the field: previous & future results

Quite remarkably, the results of this study are consistent with all the field observations and previous laboratory results of the *Artemia*-microsporidian system. Our identification of the matched and mismatched host-parasite combinations is consistent with the field data, which shows that *A. rigaudi* is dependent on its matched host to persist in the natural host community, and suggests that the same may be true for *E. artemiae* (Lievens et al. subm.). As *E. artemiae* and *A. rigaudi* performed equally poorly in their mismatched hosts, this experiment supports that suggestion. Our results for infectivity also reflect the consistently higher prevalence of *A. rigaudi* and *E. artemiae* in respectively *A. parthenogenetica* and *A. franciscana* (Lievens et al. subm., Rode et al. 2013c). In addition, we find that *A. rigaudi* is considerably more virulent than *E. artemiae* in both host species. Rode et al. (2013c) reached a similar conclusion based on the reproductive state of females collected from the field. Interestingly, the effect found by Rode et al. was that sexually mature females of both species were less likely to be brooding a clutch when they were infected with *A. rigaudi*, while in our study *A. rigaudi* did not affect the frequency of clutching once sexual maturity had been reached (data not shown). The different conditions in the field may be responsible for this seemingly additional virulence (e.g. food limitation, Brown et al. 2000, Bedhomme et al. 2004, Vale et al. 2011; temperature, Mitchell et al. 2005, Vale et al. 2008).

Further insights into the relationship between the microsporidians and their *Artemia* hosts could come from experimental coinfections. So far, we have examined the effects of *A. rigaudi* and *E. artemiae* in isolation, but coinfections are very common in the field (Lievens et al. subm.). Coinfection often has profound effects on the expression of parasite virulence and the success of their transmission, and can thus be expected to affect the evolution of microsporidian life history and host responses (Rigaud et al. 2010, Alizon et al. 2013). Studying the effects of single vs. mixed infections could therefore provide new perspectives into selection on ecological specialization in the field.

## Conclusion

In nature, multi-host parasites and multi-parasite hosts are likely to be the rule, rather than the exception (Cleaveland et al. 2001, Taylor et al. 2001, Streicker et al. 2013). Despite important research efforts in these complex systems, we still know little about the interplay between parasite specialization and its component traits (Rutrecht and Brown 2009, Rigaud et al. 2010, Hall et al. 2017). In this study, we dissected the fitness traits involved in parasite adaptation in all combinations of a naturally occurring two-host, two-parasite system. We showed that both parasites are partially specialized, with each performing better on one of the two host species. Furthermore, studying the underlying life-history traits revealed that the heart of this specialization is the delicate balance between over- and under-exploitation of the host: the drivers of infection success were spore production and the ‘tuning’ of parasite virulence to match it. This occurred despite the ecological and phylogenetic similarity of the hosts and parasites, highlighting the difficulty of adapting (or not) to multiple host species.

## Acknowledgements

We warmly thank C. Gilliot and R. Zahab for their help running the experiment and with PCR testing, T. Aubier for his assistance with the transmission assay, R. Blatrix and J.-P. Brizard for their help with the spore counting protocol, D. Degueldre for help preparing the experimental material, and the CNRS security team for regulating the light cycle on weekends. We are also grateful to A. B. Duncan, M. A. Duffy, and E. Decaestecker for constructive comments on the manuscript. YM and TL acknowledge support from CNRS and IRD.

## Author contributions

Conceptualization, E.J.P.L., Y.M. and T.L.; Methodology, P.A., J.P. and E.J.P.L.; Investigation, J.P. and E.J.P.L.; Formal Analysis, E.J.P.L. and J.P.; Writing – Original Draft, E.J.P.L.; Writing – Review and Editing, E.J.P.L., Y.M. and T.L.; Supervision, Y.M. and T.L.

## Supplementary material

**Supplementary Table 1.** Host survival during the infectivity experiment.

| Parasite species | Dose (spores/individual) | Nb. exposed | Nb. died | % Survived |
| --- | --- | --- | --- | --- |
| ***A. franciscana*** |  |  |  |  |
| Controls | 0 | 17 | 7 | 59% |
| *A. rigaudi* | 400 | 20 | 3 | 85% |
|  | 800 | 20 | 3 | 85% |
|  | 1 600 | 20 | 1 | 95% |
|  | 3 200 | 20 | 6 | 70% |
|  | 6 400 | 20 | 4 | 80% |
| *E. artemiae* | 400 | 20 | 10 | 50% |
|  | 800 | 20 | 4 | 80% |
|  | 1 600 | 20 | 9 | 55% |
|  | 3 200 | 20 | 1 | 95% |
|  | 6 400 | 20 | 8 | 60% |
| ***A. parthenogenetica*** |  |  |  |  |
| Controls | 0 | 4 | 0 | 100% |
| *A. rigaudi* | 400 | 16 | 0 | 100% |
|  | 800 | 20 | 1 | 95% |
|  | 1 600 | 20 | 0 | 100% |
|  | 3 200 | 8 | 0 | 100% |
|  | 6 400 | 4 | 0 | 100% |
| *E. artemiae* | 400 | 20 | 2 | 90% |
|  | 800 | 20 | 0 | 100% |
|  | 1 600 | 20 | 1 | 95% |
|  | 3 200 | 20 | 2 | 90% |
|  | 6 400 | 20 | 2 | 90% |

**Supplementary Table 2.** Results of paired t-tests comparing host growth before and after day 30 (all treatments combined).

| Hosts | Mean difference | p (mean difference ≠ 0) |
| --- | --- | --- |
| ***A. franciscana* males** |  |  |
| Growth between days 1 & 30 | 2.1 | <0.0001 |
| Growth between days 30 & 60 | 0.0 | 0.87 |
| ***A. franciscana* females** |  |  |
| Growth between days 1 & 30 | 2.8 | <0.0001 |
| Growth between days 30 & 60 | 0.3 | <0.0001 |
| ***A. parthenogenetica*** |  |  |
| Growth between days 1 & 30 | 1.7 | <0.0001 |
| Growth between days 30 & 60 | 0.3 | <0.0001 |

**Supplementary Table 3.**
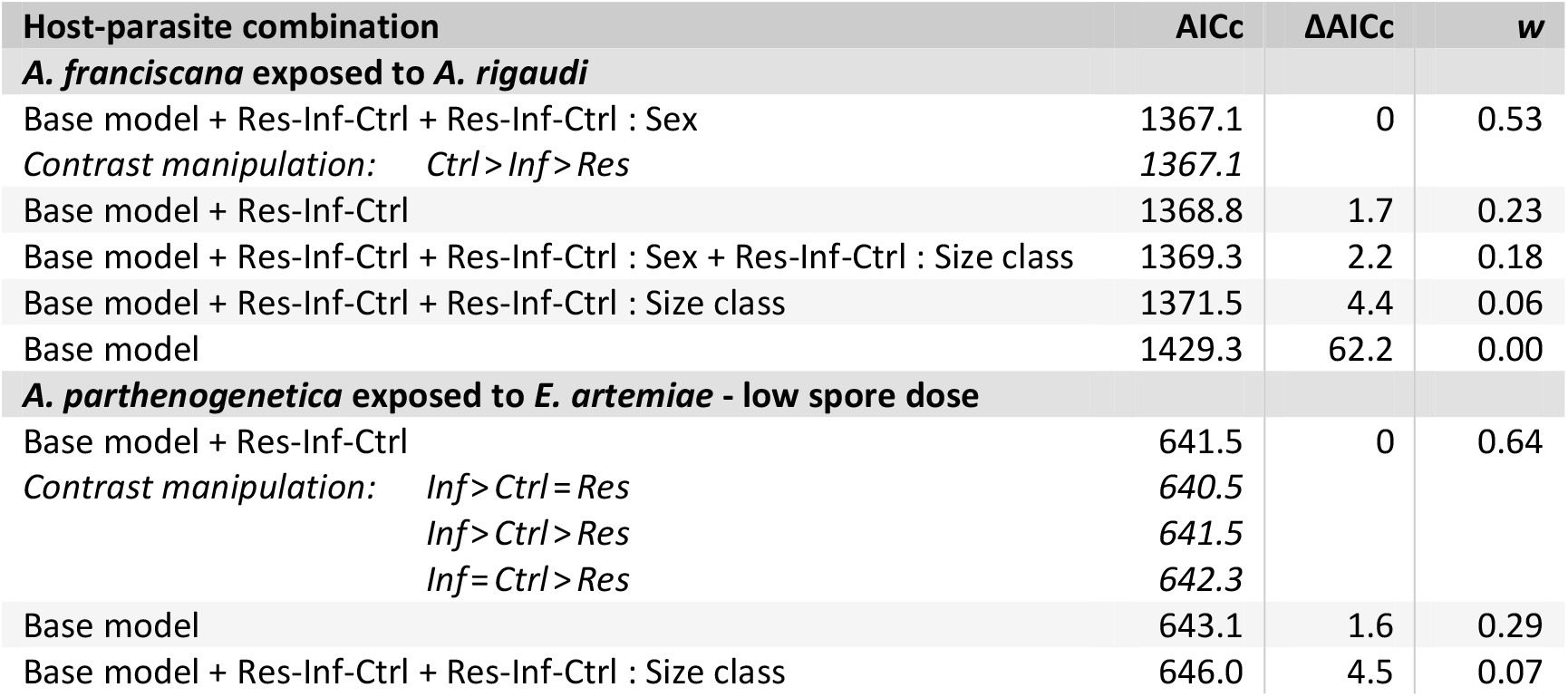
Model comparison: link between survival and infection success. For each host-parasite combination, these models grouped all the individuals exposed to that parasite (possible outcomes: uninfected or infected) and the control individuals for that host into the factor *Resistant-Infected-Control* (abbreviated Res-Inf-Ctrl). *Resistant-Infected-Control* was allowed to interact with all of the experimentally manipulated factors (the base model). The base model for *A. franciscana* included *Sex*\**Size class* and a frailty component for *Origin* (lognormal distribution, see Results). The base model for *A. parthenogenetica* included *Size class* and a frailty component for *Batch* (log-logistic distribution, see Results). We used contrast manipulation to detect how resistant, infected and control individuals differed (only models within ΔAICc = 3 of the best contrast-manipulated model are shown). Note that these analyses only included individuals that survived until at least day 15, when infection status could be definitively determined. *w* is the Akaike weight of each model.

| Host-parasite combination | AICc | $\Delta AICc$ | *w* |
| --- | --- | --- | --- |
| ***A. franciscana* exposed to *A. rigaudi*** |  |  |  |
| Base model + Res-Inf-Ctrl + Res-Inf-Ctrl : Sex | 1367.1 | 0 | 0.53 |
| Contrast manipulation: Ctrl>Inf>Res | 1367.1 |  |  |
| Base model + Res-Inf-Ctrl | 1368.8 | 1.7 | 0.23 |
| Base model + Res-Inf-Ctrl + Res-Inf-Ctrl : Sex + Res-Inf-Ctrl : Size class | 1369.3 | 2.2 | 0.18 |
| Base model + Res-Inf-Ctrl + Res-Inf-Ctrl : Size class | 1371.5 | 4.4 | 0.06 |
| Base model | 1429.3 | 62.2 | 0.00 |
| ***A. parthenogenetica* exposed to *E. artemiae* - low spore dose** |  |  |  |
| Base model + Res-Inf-Ctrl | 641.5 | 0 | 0.64 |
| Contrast manipulation: Inf>Ctrl=Res | 640.5 |  |  |
| Inf>Ctrl>Res | 641.5 |  |  |
| Inf=Ctrl>Res | 642.3 |  |  |
| Base model | 643.1 | 1.6 | 0.29 |
| Base model + Res-Inf-Ctrl + Res-Inf-Ctrl : Size class | 646.0 | 4.5 | 0.07 |

**Supplementary Table 4.** Model comparison: link between reproduction and infection success. For each host-parasite combination, these models grouped all the individuals exposed to that parasite (possible outcomes: uninfected or infected) and the control individuals for that host into the factor *Resistant-Infected-Control* (abbreviated Res-Inf-Ctrl). *Resistant-Infected-Control* was allowed to interact with all of the experimentally manipulated factors (the base model). The base model for *A. franciscana* included *Size class* and *Origin* as a random or frailty component; the base model for *A. parthenogenetica* included *Size class* and *Batch* as a random effect. We used contrast manipulation to detect how resistant, infected and control individuals differed (only models within ΔAICc = 3 of the best contrast-manipulated model are shown). Note that these analyses only included individuals that survived until at least day 15, when infection status could be definitively determined. *w* is the Akaike weight of each model.

| Host-parasite combination | AICc | ΔAICc | w |
| --- | --- | --- | --- |
| **A. franciscana exposed to A. rigaudi: time until sexual maturity** |  |  |  |
| Base model + Res-Inf-Ctrl | 951.6 | 0 | 0.82 |
| Contrast manipulation: *Inf = Res > Ctrl* | 949.7 |  |  |
| *Inf > Res > Ctrl* | 951.8 |  |  |
| *Inf > Res = Inf* | 953.2 |  |  |
| Base model + Res-Inf-Ctrl + Res-Inf-Ctrl : Size class | 954.6 | 3.0 | 0.18 |
| Base model | 974.4 | 22.8 | 0.00 |
| **A. franciscana exposed to A. rigaudi: probability of reproduction†** |  |  |  |
| Base model + Res-Inf-Ctrl | 205.5 | 0 | 0.75 |
| Contrast manipulation: *Ctrl > Inf = Res* | 204.0 |  |  |
| *Ctrl > Inf > Res* | 205.5 |  |  |
| Base model + Res-Inf-Ctrl + Res-Inf-Ctrl : Size class | 207.8 | 2.3 | 0.24 |
| Base model | 239.9 | 34.4 | 0.00 |
| **A. franciscana exposed to A. rigaudi: clutch type (higher = more nauplii)†** |  |  |  |
| Base model + Res-Inf-Ctrl | 306.7 | 0 | 1.00 |
| Contrast manipulation: *Res > Inf > Ctrl* | 306.7 |  |  |
| *Res = Inf > Ctrl* | 307.3 |  |  |
| Base model | 326.4 | 19.7 | 0.00 |
| **A. franciscana exposed to A. rigaudi: rate of offspring production\*†** |  |  |  |
| Base model + Res-Inf-Ctrl | 96.0 | 0 | 0.66 |
| Contrast manipulation: *Res > Ctrl = Inf* | 95.4 |  |  |
| *Res > Ctrl > Inf* | 96.0 |  |  |
| *Res = Ctrl > Inf* | 98.2 |  |  |
| Base model | 98.3 | 2.3 | 0.21 |
| Base model + Res-Inf-Ctrl + Res-Inf-Ctrl : Size class | 99.3 | 3.3 | 0.13 |
| **A. franciscana exposed to A. rigaudi: Fitness (LRS)\*†** |  |  |  |
| Base model + Res-Inf-Ctrl | 1094.0 | 0 | 1.00 |
| Contrast manipulation: *Ctrl > Res = Inf* | 1095.5 |  |  |
| *Ctrl > Res > Inf* | 1097.6 |  |  |
| Base model | 1145.7 | 51.7 | 0.00 |
| Base model + Res-Inf-Ctrl + Res-Inf-Ctrl : Size class | - | - | - |
| **A. parthenogenetica exposed to E. artemiae - low spore dose: clutch type (higher = more nauplii)** |  |  |  |
| Base model + Res-Inf-Ctrl + Res-Inf-Ctrl : Size class | 315.6 | 0 | 0.76 |
| Contrast manipulation: *order of effects dependent on Size class* | 313.3 |  |  |
| *order of effects dependent on Size class* | 313.4 |  |  |
| *order of effects dependent on Size class* | 315.6 |  |  |
| Base model + Res-Inf-Ctrl | 318.3 | 2.7 | 0.20 |
| Base model | 321.2 | 5.6 | 0.05 |
| **A. parthenogenetica exposed to E. artemiae - low spore dose: rate of offspring production\*** |  |  |  |
| Base model | 181.1 | 0 | 0.96 |
| Base model + Res-Inf-Ctrl | 187.3 | 6.2 | 0.04 |
| Base model + Res-Inf-Ctrl + Res-Inf-Ctrl : Size class | 196.9 | 15.8 | 0.00 |
| **A. franciscana exposed to A. rigaudi: Fitness (LRS)\*** |  |  |  |
| Base model + Res-Inf-Ctrl | 1443.2 | 0 | 0.86 |
| Contrast manipulation: *Inf > Res = Ctrl* | 1441.4 |  |  |
| *Inf = Res > Ctrl* | 1441.8 |  |  |
| *Inf > Res > Ctrl* | 1443.2 |  |  |
| Base model + Res-Inf-Ctrl + Res-Inf-Ctrl : Size class | 1446.8 | 3.6 | 0.14 |
| Base model | 1466.5 | 23.3 | 0.00 |

\*Shown for the models that weighted nauplii and cysts equally; giving either offspring type a double weight produces qualitatively equivalent results. †Only two resistant females reproduced, so these results should be interpreted with caution.

**Supplementary Figure 1.**
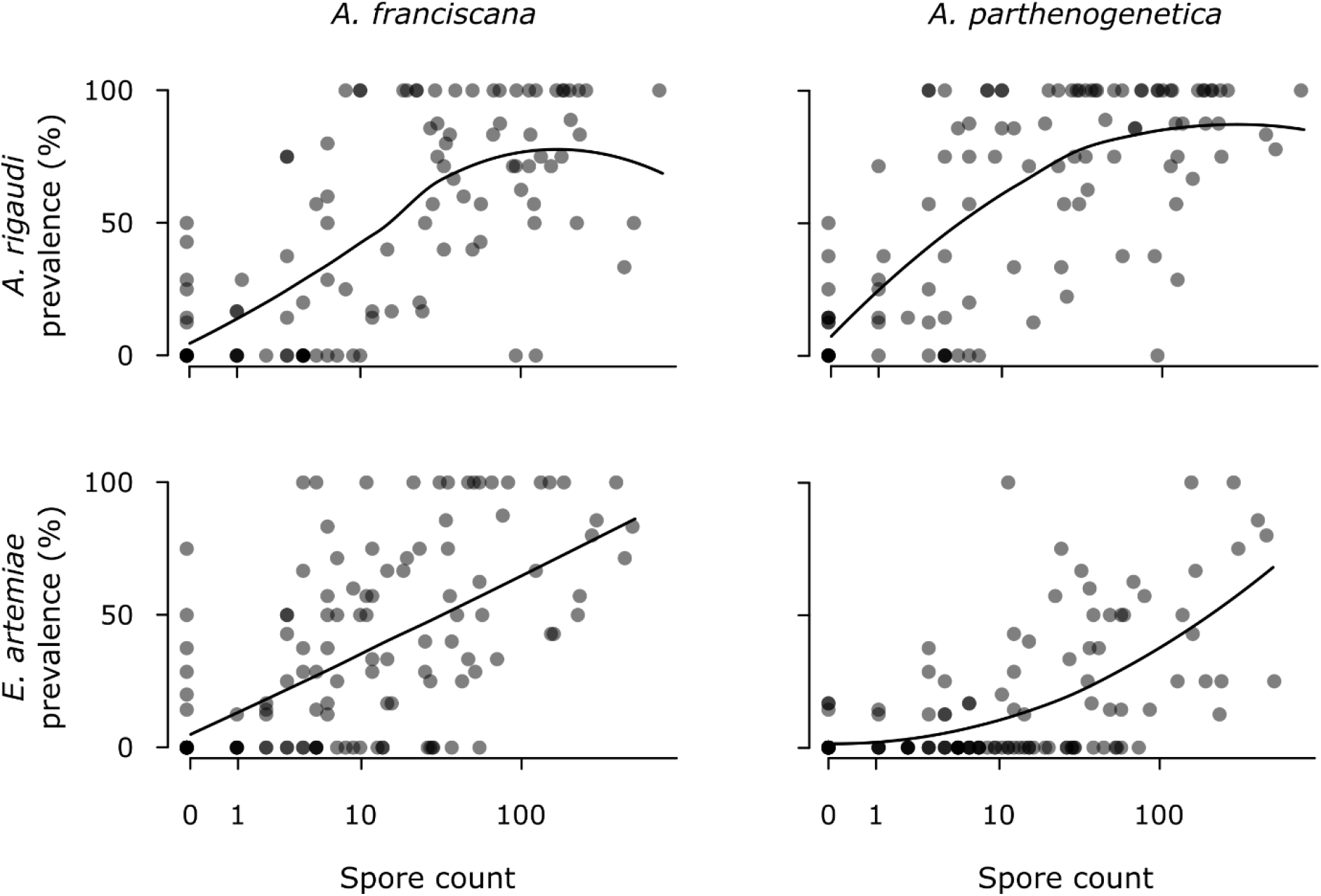
Spore production and host-to-host transmission success in the four host-parasite combinations. These graphs relate the infection success (percentage of recipients infected) to the spore count in the corresponding spore sample (*ln* + 1 scale) for *A. rigaudi* (top) and *E. artemiae* (bottom). Note that the graphs are divided by recipient species, not donor species (see Methods). Each point represents a recipient group; overlapping points shade to black. Lines represent 2^nd^-degree polynomial local regression (LOESS) fittings.

**Supplementary Figure 2.**
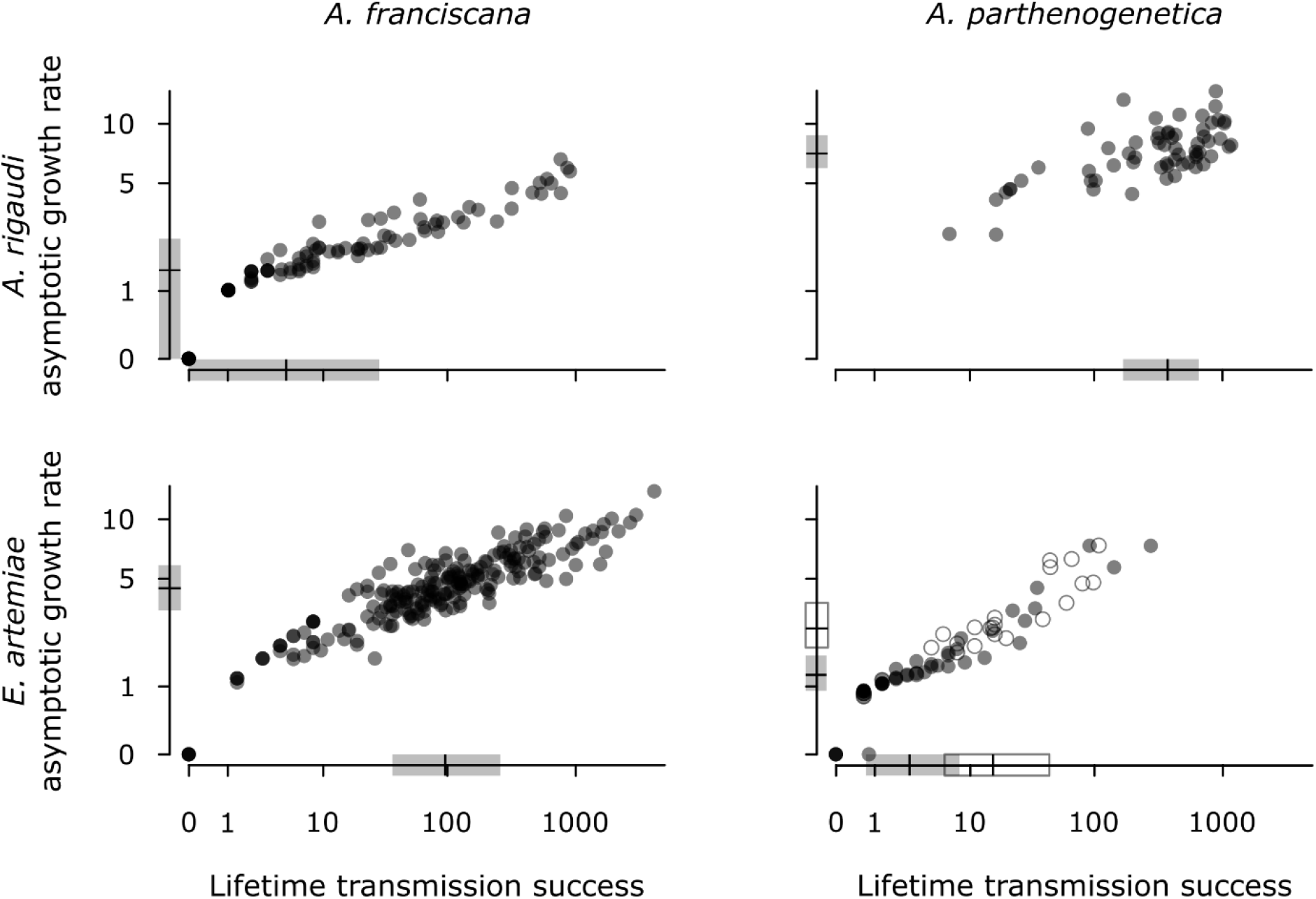
Overall fitness measures of *A. rigaudi* (top) and *E. artemiae* (bottom) infections. The asymptotic growth rate (*ln* + 1 scale) is shown as a function of the lifetime transmission success (*ln* + 1 scale). The asymptotic growth rate should be maximized during epidemics, while the lifetime transmission success, as an estimator of R0, should be maximized in endemic conditions. The median, first and third quartiles are shown by boxplots on the axes. For *A. parthenogenetica* infected with *E. artemiae*, the open circles and boxplot represent the females exposed to a high spore dose. Each point represents an infected host; overlapping points shade to black.

## Supplementary methods

### Detailed description of analyses in Table 2 (Methods > Experiment 2: Virulence and transmission > Statistical analyses: virulence & transmission)

First, we analyzed the virulence of infections (effect of the parasite on host survival, growth, reproduction, and overall fitness). This was done separately for *A. franciscana* and *A. parthenogenetica*. *A. parthenogenetica* exposed to low and high doses of *E. artemiae* were treated as separate treatments. Unless otherwise specified, our analyses proceeded as follows: we included our experimentally manipulated factors in a full regression model, used likelihood ratio tests to test their significance, and where relevant carried out post-hoc comparisons using Dunnett’s comparisons with a control (i.e. infected-with-*A. rigaudi* vs. controls, infected-with-*E. artemiae* vs. controls). Importantly, we only analyzed virulence once we could be certain of individuals’ infection status. To do this, we excluded all individuals that died before day 15 (see Methods > Experiment 2: Virulence and transmission > Experimental design and execution), and only compared infected with control individuals. To make sure that we were not missing important events occurring before this cutoff, we repeated all statistical models for exposed vs. control individuals that died before day 15.

We analyzed host survival using parametric survival models. We established a full fixed-effects model for each host species, then determined the best-fitting parametric distribution (Weibull, exponential, extreme, Gaussian, logistic, lognormal, log-logistic, Rayleigh) using the corrected AIC (Hurvich and Tsai 1989). We then tested the significance of the predictive effects as described above. Finally, we confirmed the fit of the model by performing a goodness-of-fit test (comparing the likelihood of the observed data with the likelihood distribution of simulated datasets based on the model predictions). The full model for *A. franciscana* included *Treatment*, *Sex*, *Size class*, and all double interactions. The full model for *A. parthenogenetica* included *Treatment*, *Size class*, and their interaction. *Origin* and *Batch* were included as frailty components for *A. franciscana* and *A. parthenogenetica*, respectively, as they could introduce heterogeneity in mortality rates. Data were right-censored on day 60.

To test the effects of parasite infection on growth, we first checked whether there was significant growth between days 1 & 30 and days 30 & 60 (paired t-tests of the size difference between day 30 & 1 and day 60 & 30). Most growth occurred during the first month (see Results), so we analyzed growth between days 1 & 30 further using linear mixed models. For *A. franciscana*, we looked at the effects of the fixed effects *Sex*, *Treatment*, *Size class* and all their interactions, with *Origin* as a random effect. For *A. parthenogenetica*, the full model included *Treatment*, *Size class* and their interaction as fixed effects, and *Batch* as a random effect.

To analyze (female) reproductive success, we decomposed female reproduction into a) time until sexual maturity, b) the probability of producing a clutch, c) the rate of offspring production, d) the timing of offspring production, and e) the type of offspring produced. All models included *Treatment*, *Size class*, and their interaction as fixed effects, and *Origin* or *Batch* as random (or frailty) effects. The response variables and statistical models were as follows. a) Time until sexual maturity: the number of days until females became sexually mature, analyzed using parametric survival models. As above, we first determined the best survival distributions to use, then tested the significance of the predictive effects. Females were right-censored in case of death. b) Probability of producing a clutch: a binary variable describing whether a female produced a clutch during the experiment or not, analyzed using generalized linear mixed models with a Bernouilli distribution. c) Rate of offspring production: for females that produced at least one clutch, the total number of offspring divided by the length of the reproductive period. The length of the reproductive period was defined as the difference between the date of death (or censoring) and the date of maturity. The data were analyzed using linear mixed models. Offspring could be nauplii or cysts, and these two offspring types were not directly comparable (they probably require different amounts of energy to produce, and we allowed mortality to occur before counting nauplii). To account for this, we repeated the analyses with nauplii weighted twice, equally, or half as much as cysts, and based our conclusions on the overall pattern. d) Timing of offspring production: for females that produced at least one clutch, the clutch size through time. Clutch size was modelled as a quadratic function of clutch date, with clutch date expressed as the elapsed proportion of the female’s reproductive period (e.g. for two females reproducing on the 10^th^ day of sexual maturity, where one died on the 20^th^ day and one was censored on the 40^th^, the elapsed proportions would be 0.5 and 0.25). Timing was analyzed using generalized linear mixed models with a negative binomial distribution; *Individual* was included as a random variable to control for pseudoreplication. As in (c), we ran models where nauplii were weighted twice, equally, or half as much as cysts, and based our conclusions on the overall pattern. e) Type of offspring produced: for females that produced at least one clutch, a binomial combination of the number of clutches consisting of nauplii vs. cysts, analyzed using generalized linear mixed models.

As a final virulence measure, we estimated the fitness of (female) hosts. Our fitness proxy was the lifetime reproductive success (LRS), calculated as the total number of offspring produced over the study period. This produced a zero-inflated count distribution, to which we fit negative binomial hurdle models. The full models included *Treatment*, *Size class*, and their interaction as fixed effects; random effects (such as *Origin* and *Batch*) were not supported by the package. As above, we ran models where nauplii were weighted twice, equally, and half as much as cysts, and based our conclusions on the overall pattern.

Next, we analyzed the parasites’ transmission (spore production rate, infectiousness of a single spore, and overall fitness). These analyses were combined for infections in *A. franciscana* and *A. parthenogenetica*. Unless otherwise specified, we included our experimentally manipulated factors in a full regression model, and used likelihood ratio tests to test their significance. If relevant, post-hoc comparisons were carried out using Tukey comparisons.

To estimate the infectiousness of a single spore, we used the results of the transmission assay. We assumed that the establishment of microsporidian infections follows an independent-action model with birth-death processes. This model assumes that a parasite population grows in the host until it reaches an infective threshold, at which point the infection is considered to be established (Schmid-Hempel 2011 pp. 225–6). In our assay, we considered that an infection was established when we could detect it; in other words, the infective threshold corresponded to the threshold for PCR detection (estimated at ∼1 000 spores inside the host’s body, unpublished data). In these models, the probability per spore to start an infection, *p*, is equal to 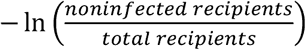 /*D* where *D* is the spore dose (Schmid-Hempel 2011 pp. 225–6). In our transmission assay, *D* can be approximated by the number of spores in the fecal sample taken from the donor at the start of the assay (= spore count transformed to spores/mL, or * 700), divided by 5*8 = 40 (fecal samples accumulated over a 5-day period but we only exposed recipients for one day; the inoculum was shared amongst 8 recipients). We calculated a value of *p* for every replicate in the transmission assay; *p* was then analyzed using linear mixed models. The model included *Recipient species*, *Parasite species*, and their interaction as fixed effects; an *Individual*-level random effect was included to control for pseudoreplication (each donor host was used to infect a group of *A. franciscana* and a group of *A. parthenogenetica* recipients; some donors were also re-used in the transmission assays on day 30 and 60).

We then tested whether the rate of spore production was dependent on the host-parasite combination. We used *Spore count*, the number of spores counted in the fecal sample, as the response variable in a generalized linear mixed model with a negative binomial distribution. We did not transform the spore count to spores/mL (≈ spore count * 700) to avoid skewing the error distribution. The fixed effects were *Host species*, *Parasite species*, and their interaction; an *Individual*-level random effect was included to control for pseudoreplication. To avoid comparing apples with oranges, we excluded *A. parthenogenetica* that had been exposed to 10000 *E. artemiae* spores from this model. However, we tested separately whether the rate of spore production differed for *A. parthenogenetica* infected with different doses of *E. artemiae* (equivalent model with fixed effect *Dose*). Spore production analyses were carried out for infected hosts only.

As a final measure of parasite success, we investigated parasite fitness in the different host-parasite combinations. For each established infection (i.e. each infected host), we used two measures of spore production as proxies for parasite fitness. First, we calculated a proxy for the ‘lifetime transmission success’: we summed the number of spores in the fecal samples taken on days 15, 30, 45 and 60 for each infection, then corrected this cumulative spore count by *p*, the average infectiousness of a single spore in a given host-parasite combination (as calculated above). Second, we calculated an asymptotic growth rate by computing the dominant eigenvalue of a standard Leslie matrix,

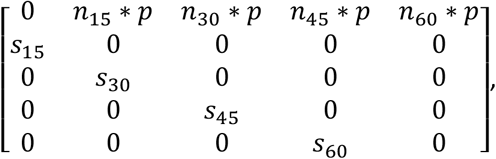

where *n_i_* is the number of spores in the fecal sample on day *i*, *p* is the average infectiousness of a single spore in that host-parasite combination (as calculated above), and *s_i_* describes whether the host survived until day *i* (1) or not (0). While the lifetime transmission success is a measure of the basic reproduction number R_0_, which describes parasite fitness under stable endemic conditions, the asymptotic growth rate is a measure of the net population growth rate, which describes fitness under epidemic conditions (Frank 1996, Hethcote 2000); we included both measures because either situation can occur in the field. We compared the two measures across host-parasite combinations using non-parametric Kruskal-Wallis tests with Dunn’s post hoc testing (R package PMCMR, Pohlert 2014). *A. parthenogenetica* exposed to low and high spore doses of *E. artemiae* were treated separately.

